# tRNA modifications tune decoding of codon pairs to prevent cellular quality control responses

**DOI:** 10.1101/2024.02.27.582385

**Authors:** Jie Wu, Cristian Eggers, Olga Sin, Łukasz Koziej, Hector Mancilla, Fabienne Mollet, Hans R. Schöler, Hannes C.A. Drexler, Tristan Ranff, Christian Fufezan, Claudine Kraft, Sebastian Glatt, Jan M. Bruder, Sebastian A. Leidel

## Abstract

tRNA modifications tune translation rates and codon optimality, thereby optimizing co-translational protein folding, but how codon optimality defects trigger cellular phenotypes remains unclear. Here, we show that ribosomes stall at specific modification-dependent codon pairs, triggering ribosome collisions and inducing a coordinated and hierarchical response of cellular quality control pathways. Ribosome profiling reveals an unexpected functional diversity for wobble-uridine (U_34_) modifications during decoding. The same modification can have different effects at the A and P sites. Furthermore, modification-dependent stalling codon pairs induce ribosome collisions, triggering ribosome-associated quality control (RQC) to prevent protein aggregation by degrading aberrant nascent peptides and mRNAs. RQC inactivation stimulates the expression of molecular chaperones to remove protein aggregates. Our results show that loss of tRNA modifications primarily disrupts translation rates of suboptimal codon pairs and reveal the coordinated regulation and adaptability of cellular surveillance systems to ensure efficient and accurate protein synthesis and maintain protein homeostasis.

## Introduction

Translation speed varies along individual mRNA transcripts thereby substantially impacting protein synthesis rates and orchestrating co-translational protein folding and protein localization^1–7^. Codon optimality is a concept that applies a supply-demand model to tRNAs and their respective codons, and best explains codon-specific translation rates. However, a major limitation is that an identical optimality value is assigned to all instances of a codon across the transcriptome, whereas it is known that numerous factors can modulate the translation elongation rate of individual codons, such as tRNA modifications, mRNA secondary structure, adjacent codons, or an amino acid’s position in nascent peptides^4,5,8–11^. Hence, it is unclear to what extent context-specific factors tune the optimality of individual codons and how this affects the quality of translation and protein synthesis.

A powerful strategy to systematically alter codon-optimality and translation dynamics is the use of tRNA modification mutants because tRNA selection depends largely on codon-anticodon interactions in the A site of translating ribosomes. While most of these interactions follow Watson-Crick rules, wobble pairing enables cells to decode multiple codons by a single tRNA. The physicochemical parameters of such base-pairing interactions are optimized by chemical modifications of nucleotides in the anticodon loop^12,13^. A classic example of this phenomenon are wobble uridines at tRNA position 34 (U_34_). U_34_ nucleotides are generally modified in eukaryotes by 5-carbamoylmethyl (ncm^5^), 5-methoxycarbonylmethyl (mcm^5^) or 5-methoxycarbonylmethyl-2-thio-uridine (mcm^5^s^2^U) through the activity of the Elongator (Elp) complex and the Urm1 pathway^14–17^. These highly conserved modifications are critical for decoding^18,19^ and their absence is associated with several human diseases, including cancers, familial dysautonomia and DREAM-PL Syndrome^20–24^. Mechanistically, loss of U_34_ modifications slows decoding of specific codons in different species^8,9,25–27^. tRNA-modification mutants can therefore be used to alter codon optimality without changing the sequence of cellular transcripts in contrast to using reporter systems. Interestingly, U_34_-modification mutants exhibit aggregation of cellular proteins, similar to what is observed in strains depleted of co-translational chaperones^9,28,29^. Under normal growth conditions, translation dynamics are optimized to balance protein folding and quality control pathways to maintain protein homeostasis^7^, and the aggregation phenotype of U_34_-modification mutants highlights the link between translation dynamics and protein quality. However, how cells respond to codon-specific translation defects and how these trigger cellular quality control systems remains poorly understood.

Perturbations in translation dynamics due to alterations of the translation machinery, defective mRNA, or environmental stress can stall ribosomes and induce ribosome collisions. Collided ribosomes are resolved by ribosome-associated quality control (RQC) and induce other cellular pathways, such as the integrated stress response (ISR) to reduce initiation rates avoiding further collisions, and no-go decay (NGD) to degrade the potentially defective mRNA^30–35^. To activate RQC, the E3-ubiquitin ligase Hel2 recognizes the extended interface formed by collided ribosomes^33,36–39^. Previous studies have shown that strong perturbations of translation, such as UV, drug-induced stress or artificial reporter constructs, induce ribosome collisions and trigger different branches of cellular quality control^35^. In contrast, codon-specific slowdowns in tRNA modification mutants are comparably subtle and much closer to physiological conditions. Hence, it is not clear whether they are sufficient to trigger the RQC pathway.

To determine how sequence context tunes the optimality of individual codons across the transcriptome and how their modulation affects cellular quality control, we applied high-resolution ribosome profiling and disome sequencing in baker’s yeast devoid of U_34_ modifications (Fig. 1a)^40–44^. We found that the previously underappreciated ncm^5^U and mcm^5^U modifications are required for wobble decoding at the ribosomal A site. Surprisingly, ncm^5^U plays a strikingly different role at the P site: Loss of this modification exacerbates slow A-site decoding when the P site contains a codon cognate to ncm^5^-modified tRNAs. Strikingly, although most suboptimal codons are translated without detectable effect, specific U_34_-dependent codon pairs induce ribosomal collisions and trigger RQC. Activation of RQC by Hel2 leads to the degradation of mRNAs enriched in difficult-to-translate codons and codon pairs, thereby contributing to the maintenance of cellular protein homeostasis as a first line of defense. RQC targets are less likely to encode aggregation-prone proteins and are enriched in suboptimal codon pairs. Importantly, when RQC is compromised, cells upregulate molecular chaperones via Hsf1 and are better able to cope with proteotoxic stress. Our findings demonstrate that codon optimality is context-dependent and that RQC coordinates with the cellular chaperone machinery in a hierarchical manner to maintain cellular protein homeostasis in response to codon-specific translation defects.

**Fig. 1.**
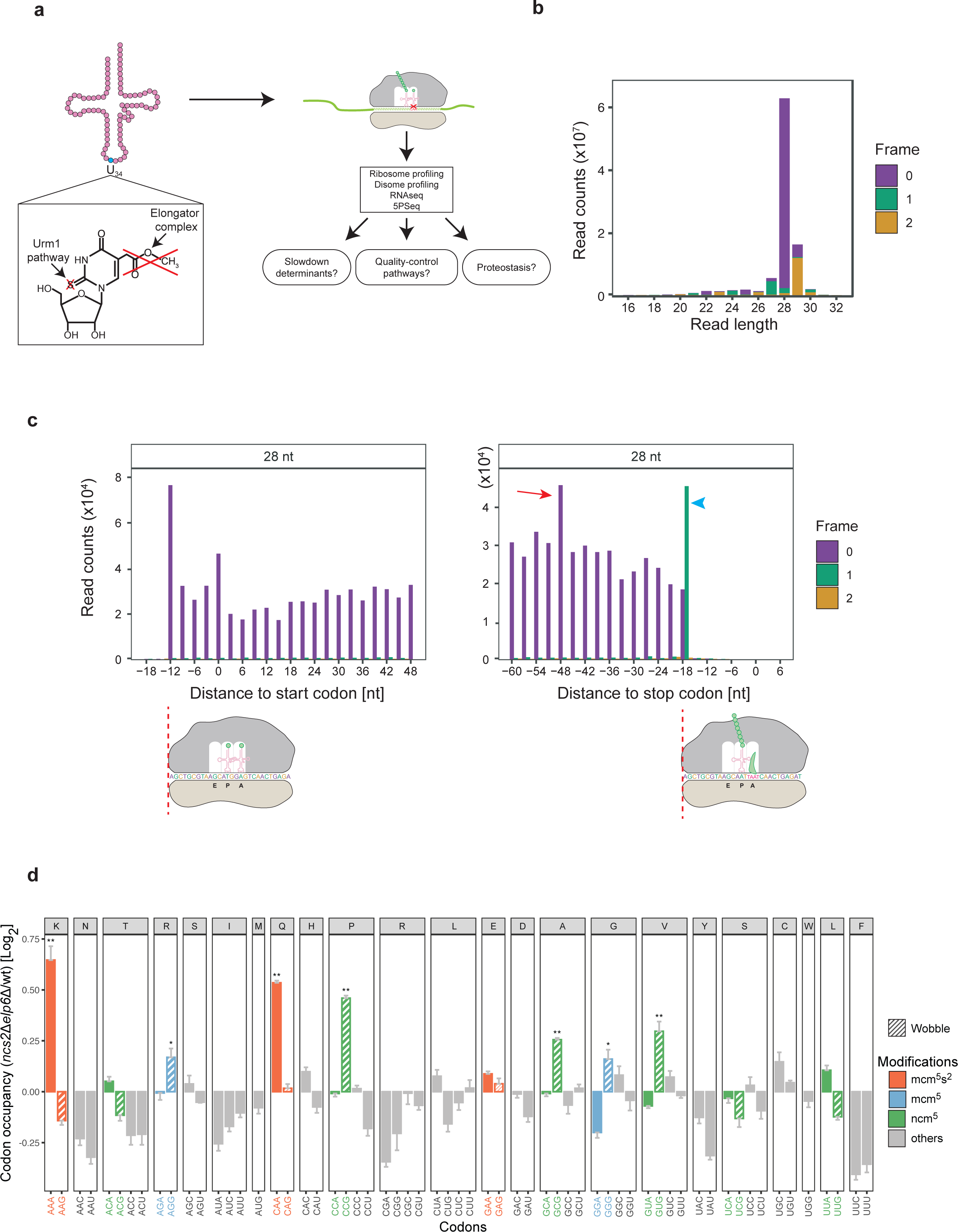
U_34_ modifications affect translation differentially. (a) Overview of this study. 5-methoxycarbonylmethyl-2-thiouridine (mcm^5^s^2^U; chemical formula shown), 5-methoxycarbonylmethyluridine (mcm^5^U), and 5-carbamoylmethyluridine (ncm^5^U) are located at wobble uridine (U_34_). Deleting *NCS2* (Urm1 pathway) and *ELP6* (Elongator complex) completely removes these modifications. Ribosome profiling, disome profiling, RNAseq, and 5PSeq were performed. (b) The majority of monosome footprints were 28 nt long and predominantly in reading frame 0. (c) The distribution of the 5’ end of 28-nt footprints is shown at the start (left) and stop (right) codon; the cartoon below depicts initiating and terminating ribosomes. The terminating ribosomes accommodate four nucleotides in the ribosomal A site due to eRF1 recognition, which is visible as an apparent frameshift at the stop codon (blue arrowhead). A peak 48 nt upstream of the stop codon indicates ribosomes that collide with terminating ribosomes (red arrow). (d) Alterations of codon occupancy in the absence of U_34_ tRNA modification. Relative A-site occupancy of each codon in *ncs2*Δ*elp6*Δ mutant cells compared to wild type. Log_2_ fold change > 0 indicates that codons are decoded slow in the mutant. Modification-dependent codons are highlighted by color: mcm^5^s^2^U (red); mcm^5^U (blue); ncm^5^U (green). Significantly slow codons with Log_2_ fold change > 0.1 are highlighted with asterisks (one-sided Welch’s t-test; \**p*-value < 0.05, \*\**p*-value < 0.01).

## Results

### U_34_ tRNA Modifications Differentially Affect Cognate and Wobble Decoding

Codon-specific translation defects have mostly been characterized globally and not at the level of individual instances throughout the transcriptome, precluding a full understanding of their context dependency. To understand the impact of U_34_ modifications on individual transcripts, we optimized our ribosome profiling protocol for improved length and reading frame of the resultant footprints, allowing us to map translation dynamics at high resolution (Fig. 1b)^40,41,43,44^. To verify the quality of our data, we aligned reads at the start and stop codon, showing the expected 3-nt periodicity pattern reflecting ribosomal movements along mRNA (Fig. 1c and Supplementary Fig. 1a and 1b). Furthermore, we observed a peak 48 nt upstream of the stop codon indicative of ribosomal queueing upstream of the terminating ribosome (48 nt=28+2+18 nt) (Fig. 1c right; red arrow). Interestingly, the footprints of terminating ribosomes (at approximately -18 nt relative to the stop codon) shift to a different reading frame within the same footprint length, which indicates that the ribosomal A site accommodates 4 nucleotides during eRF1 recognition (Fig. 1c and Supplementary Fig. 1a right; blue arrowhead)^45–49^. These examples demonstrate that our optimized libraries capture subtle physiological translation events.

The absence of mcm^5^s^2^U in 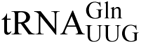 and 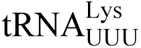 slows down the decoding of their respective cognate CAA and AAA codons^8,9^. Even though ncm^5^U and mcm^5^U are found in 8 additional tRNA and their absence leads to stress sensitivity^50^, no codon-specific translation defects have been linked to the loss of these tRNA modifications. To obtain a more granular picture, we generated a high-resolution dataset for *ncs2*Δ*elp6*Δ mutants lacking all three modifications to evaluate changes in translation speed with high precision. We used two independent normalization methods (position-based and footprint-based; see Methods) to determine changes in codon-specific translation rates. Consistent with previous work, AAA and CAA are the most affected codons using both methods (Fig. 1d and Supplementary Fig. 1c)^8,9^, and we did not observe an effect on the translation speed of codons cognate to mcm^5^U-modified tRNAs (AGA and GGA) or ncm^5^U-modified tRNAs (ACA, CCA, GCA, GUA, UCA, and UUA). Interestingly however, some of their synonymous G-ending codons (mcm^5^U: AGG and GGG; ncm^5^U: CCG, GCG, and GUG) slowed down significantly. The slow G-ending codons CCG and GCG do not have cognate tRNAs and therefore rely on wobble decoding (Supplementary Fig. 1d)^51^. While the remaining slow G-ending codons AGG, GGG, and GUG have cognate tRNAs, these tRNAs are not essential in yeast, demonstrating that these codons rely at least in part on wobble decoding by essential tRNAs^50^. In contrast, none of the codons synonymous to the mcm^5^s^2^U-modified tRNA (AAG, CAG and GAG) are significantly slowed down, demonstrating that mcm^5^s^2^U behaves differently from mcm^5^U and ncm^5^U. Together, we show that the previously underappreciated ncm^5^U and mcm^5^U modifications are required for U**•**G wobble pairing but not for cognate decoding in the ribosomal A site (Fig. 1d and Supplementary Fig. 1c and 1d).

A global slowdown in the decoding of specific codons could be caused by the arrest of ribosomes in a specific state during the elongation cycle^26,52^. Therefore, we used single-particle cryo-EM to compare the overall distribution of ribosomal states in the wild type and *ncs2*Δ*elp6*Δ cells. To this end, we isolated monomeric 80S ribosomes using a sucrose gradient after nuclease digestion and collected two independent datasets for each of the strains (Supplementary Table 1). From these images, we were able to reconstruct individual structures of translating ribosomes with an overall resolution of up to 1.8 Å and distinguish six distinct elongation states (Supplementary Fig. 2a-e). However, we did not observe major changes in the distribution of elongation states between the two strains, except for a marginal enrichment of post-translocation (POST) states detectable in the strain lacking U_34_ modifications (*ncs2*Δ*elp6*Δ; Supplementary Fig. 2b). These results are consistent with our ribosome profiling data, indicating less efficient decoding of certain codons by unmodified tRNA in the A site of ribosomes. The results also suggest that the observed phenotypic changes are caused by defects in specific decoding events and that the lack of U_34_ modification does not generally trap translating ribosomes in a specific state. Therefore, we sought to understand whether specific translation events underlie the observed translational slowdown.

### tRNA Modifications are Critical for the Decoding of Codon Pairs

The translation speed of codons within the same transcript can vary substantially^40,53,54^. However, codon optimality is generally assigned to codons regardless of their position in the transcriptome. The high resolution of our dataset allowed us to investigate whether the sequence context affects the optimality of individual instances of a codon. Therefore, we first asked which codons in the transcriptome are most affected in modification mutants. We computed a vulnerability score for each codon by comparing the ratio of pause scores between *ncs2*Δ*elp6*Δ mutants (lacking the ncm^5^U, mcm^5^U, and mcm^5^s^2^U modifications) and wild-type yeast, which quantifies how sensitive a particular codon is to the absence of these tRNA modifications (see Methods). While AAA and CAA codons are on average decoded more slowly in *ncs2*Δ*elp6*Δ cells (Fig. 1d and Supplementary Fig. 1c), the extent of the slowdown varies significantly for different instances of the same codon, even within the same transcript (Fig. 2a). We ranked all codons in the transcriptome based on their vulnerability scores and divided them into 10 equally sized bins from low to high vulnerability (Fig. 2b). Strikingly, codons that depend on U_34_ modifications (AAA, CAA, AGG, GGG, CCG, GCG, and GUG) are enriched in high vulnerability bins compared to low vulnerability bins (Fig. 2b bottom), while modification-unrelated codons are evenly distributed, demonstrating that our method is suitable for identifying and ranking codons according to their sensitivity to the absence of U_34_ modifications.

**Fig. 2.**
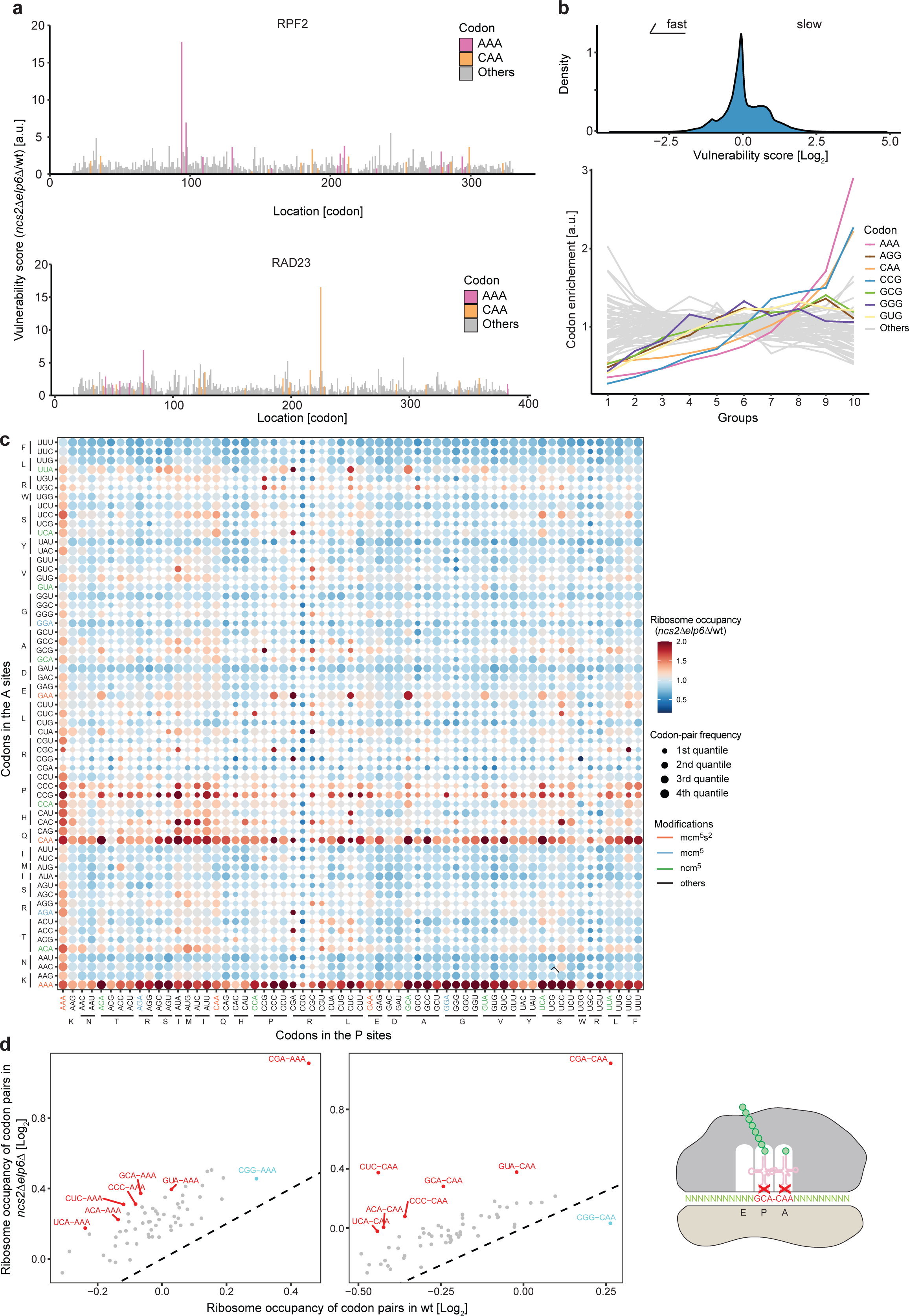
Strong pausing occurs at tRNA modification-dependent codon pairs. (a) The distribution of vulnerability scores for each codon in two example genes, *RPF2* and *RAD23*. Vulnerability scores are calculated by normalizing the pause score of codons in mutant yeast to those in wild type. AAA (magenta) and CAA (orange) are highlighted. 15 codons at the start and end of the coding sequence are excluded from the calculation. (b) The distribution of vulnerability scores from all codons in the transcriptome (top). Codons are equally divided to 10 groups according to the rank of vulnerability scores. The enrichment of each codon in each group is calculated. U_34_-modification-dependent codons (according to Fig. 1d; marked by colors) are enriched in the groups with higher vulnerability scores. (c) Two-dimensional codon plot depicting the change of ribosome occupancy for codon pairs between the *ncs2*Δ*elp6*Δ mutant and wild type. Indicated on the x axis are the codons in the ribosomal P site; on the y axis are the codons in the ribosomal A site. Dark red color indicates a high ratio of ribosome occupancy in the *ncs2*Δ*elp6*Δ mutant compared to wild type. For better visualization, ratios ≥ 2 were set to 2. Size of the dots represents the frequency of each codon pair in yeast categorized into four quantiles. Codons cognate to U_34_ modified tRNAs are indicated by color: mcm^5^s^2^U (red); mcm^5^U (blue); ncm^5^U (green). (d) Specific codon pairs are selected from (c). Ribosome occupancy between *ncs2*Δ*elp6*Δ and wild-type yeast were compared for NNN-AAA (left) and NNN-CAA (right) codon pairs. The diagonal line indicates no change between mutant and wild type. Codon pairs that are found more frequently than the other pairs are highlighted in red. CGG-AAA and CGG-CAA (highlighted in blue) were weakly or not affected, and CGG-CAA was used as a negative control in Fig. 3. The diagram below depicts a pausing ribosome translating a pair of U_34_-modification-dependent codons in the P and A sites, such as GCA-CAA.

Next, we used the highly vulnerable codons and searched for features that can explain why they are more affected by a lack of tRNA modifications than less vulnerable codons. Several studies have indicated that the region close to the 5’ end of transcripts is enriched with nonoptimal codons^55,56^. However, we did not observe an enrichment of vulnerable codons in specific regions of the ORF including the 5’ end (Fig. 2a and Supplementary Fig. 3a). Instead, we found that codons encoding for the negatively charged aspartic acid and glutamic acid are depleted in the P site of vulnerable AAA, CAA, and GAA codons but not U_34_-independent codons such as UAU suggesting that electrostatic interactions of the amino acids in the P site reduce the requirement of U_34_ tRNA modifications for A-site decoding (Supplementary Fig. 3b). However, the amino acids enriched in the P site do not share chemical properties and therefore did not appear to explain codon vulnerability.

Therefore, we asked whether the extent of slowdown is modulated by the codon context. To visualize synergistic effects of adjacent codons in the *ncs2*Δ*elp6*Δ mutant, we generated a 2-dimensional plot depicting the change of translation speed at the P-A sites compared to wild type (Fig. 2c). As expected, AAA, CAA and other U_34_-dependent codons are overall slow during A-site decoding independent of the P-site codon, which is indicated by horizontal red lines in the plot. Interestingly, we observed several strong stalling codon pairs when CGA is located in the P site and a codon decoded by an U_34_-modified tRNA is in the A site, such as CGA-AGA, CGA-CAA (Supplementary Fig. 3b). This is consistent with the observation that P-A codon pairs affect translation efficiency^10,57^, and suggests that CGA in the P site enhances the dependence on U_34_ modifications for the decoding of A-site codons, likely by altering the secondary structure of the mRNA around the decoding center^57^. Surprisingly, codons cognate to ncm^5^U-modified tRNA (ACA, GCA, GUA, and UCA) at the P site slow down A-site decoding of mcm^5^s^2^U-dependent codons (AAA, CAA, and GAA) in *ncs2*11*elp6*11 yeast (Fig. 2d and Fig. 3d and 3e) while these ncm^5^U-modified codons are decoded at a normal pace in the ribosomal A site (Fig. 1d and Supplementary Fig. 1c). These data show that hypomodified tRNAs in the P site synergistically enhance the requirement of modifications for decoding A-site codons and that codon vulnerability can be largely explained by codon pairs. Finally, U_34_ modifications play strikingly different roles at the A site and at the P site: while a lack of ncm^5^U affects wobble decoding at the A site, ncm^5^U deficiency impairs the cognate codons at the P site. From this we conclude that U_34_ modifications on tRNAs at both ribosomal sites are critical for tuning translation elongation.

**Fig. 3.**
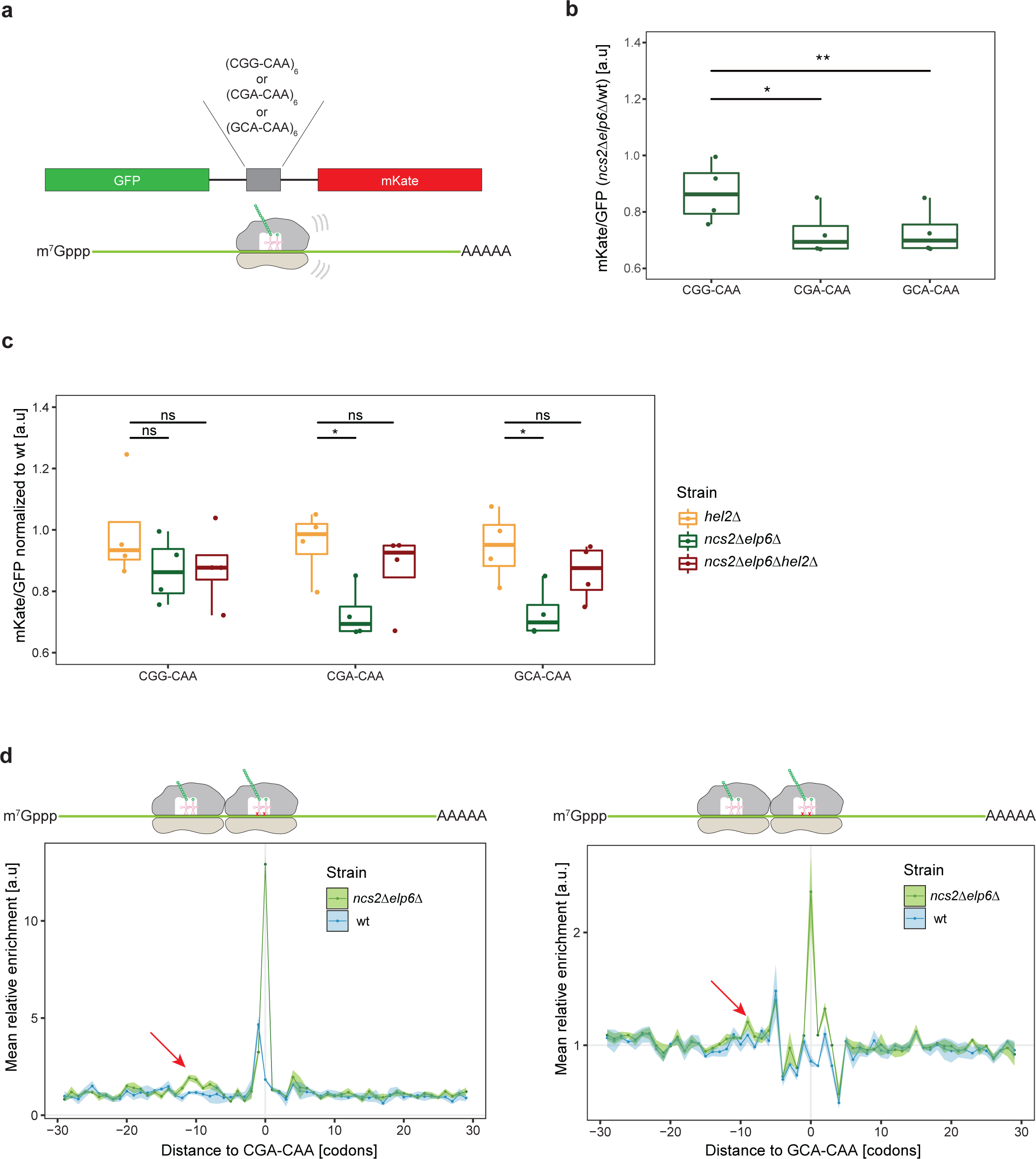
Ribosomes stall on slow codon pairs. (a) Schematic of the stalling reporter: 6 copies of CGA-CAA (slow), GCA-CAA (slow), or CGG-CAA (fast) were inserted between GFP and mKate. (b) The mKate/GFP ratios of CGG-CAA, CGA-CAA and GCA-CAA reporters in *ncs2*11*elp6*11 strains were normalized to wild type. Both stalling pairs show a decreased ratio in comparison to the control pair (two-sided student’s t-test; \**p*-value < 0.05, \*\**p*-value < 0.01). (c) Similar to (b) for *hel2*Δ, *ncs2*Δ*elp6*Δ and *ncs2*Δ*elp6*Δ*hel2*Δ mutants. Knocking out *HEL2* in addition to *NCS2* and *ELP6* rescues the stalling effects of codon pairs. A *HEL2* knockout was used as a control (two-sided student’s t-test; \**p*-value < 0.05). (d) The distribution of ribosome occupancy (monosomes) around CGA-CAA (left) and GCA-CAA (right). Colliding ribosomes ∼10 codons upstream of the codon pairs in the *ncs2*Δ*elp6*Δ mutant are highlighted by red arrows. The shaded area indicates the degree of experimental variation within three replicates. The cartoon above depicts collided ribosomes relative to the plot.

### tRNA Modification Defects Induce Ribosome Collision

To further substantiate that loss of tRNA modifications induces ribosomal stalling at U_34_-dependent codon pairs, we generated reporters containing six consecutive codon pairs encoded between GFP and mKate (Fig. 3a)^58^. For this purpose, we combined the slow mcm^5^s^2^U-dependent codon CAA with CGA or the ncm^5^U-dependent GCA, or used six consecutive CGG-CAA pairs as a negative control (Fig. 3a). When ribosomes stall during translation of the non-optimal codon pairs, less mKate is synthesized, as indicated by a low mKate/GFP ratio. Indeed, both slow codon pairs showed a significant decrease of ∼20 % in the mKate/GFP ratio when comparing wild type to the *ncs2*Δ*elp6*Δ mutant, suggesting that ribosomes stall at U_34_-dependent non-optimal codon pairs (Fig. 3b). Strong stalling induces ribosomal collisions and triggers the RQC pathway^37^. Therefore, we asked whether RQC is triggered in *ncs2*Δ*elp6*Δ cells expressing the stalling reporters and analyzed a *ncs2*Δ*elp6*Δ*hel2*Δ mutant lacking Hel2, the E3- ubiquitin ligase that senses collided ribosomes upstream of the RQC pathway^33,37,39,59^. Indeed, deleting *HEL2* increases the mKate/GFP ratio for the non-optimal CGA-CAA and GCA-CAA codon pairs, but not for the CGG-CAA control (Fig. 3c), confirming that RQC is activated at U_34_-modification-dependent codon pairs.

To verify that ribosomes similarly stall on endogenous transcripts, we searched our ribosome profiling data for evidence of colliding ribosomes, characterized by a peak in ribosome occupancy 10 codons upstream of the A site of the stalled ribosomes^60^. Strikingly, we observed such a peak ∼10 codons upstream of CGA-CAA (Fig. 3d left), GCA-CAA (Fig. 3d right) and CGA-AAA pairs (Supplementary Fig. 4a) while a similar peak is not found upstream of single AAA or CAA codons (Supplementary Fig. 4b), indicating that ribosome collisions are induced by stalling at tRNA modification-dependent codon pairs rather than at single codons. Upon recognition of collided ribosomes, Hel2 ubiquitinates uS10 and synthesizes ubiquitin chains on uS3 downstream of Mag2 at specific lysine residues^61–63^. To test whether RQC is active in U_34_-modification mutants we assessed the ubiquitination of disomes from *ncs2*Δ*elp6*Δ and *ncs2*Δ*elp6*Δ*hel2*Δ cells by mass spectrometry and observed K212 ubiquitination of uS3 and K6/8 ubiquitination of uS10 in *ncs2*Δ*elp6*Δ cells. However, uS10 ubiquitination was not detectable on ribosomes from *ncs2*Δ*elp6*Δ*hel2*Δ cells, and uS3 ubiquitination levels were diminished (Supplementary Fig. 5a). These findings demonstrate that specific U_34_-dependent codon pairs induce strong ribosomal stalling in tRNA modification mutants *in vivo*, leading to ribosome collisions that trigger the RQC pathway via Hel2.

### The RQC and ISR pathways remedy codon-specific translation defects

Collided ribosomes can be specifically studied by extracting disomes instead of monosomes during ribosome profiling^42,64–67^. To more specifically analyze ribosome collisions at U_34_-dependent non-optimal codon pairs of endogenous transcripts, we generated disome libraries of *hel2*Δ, *ncs2*Δ*elp6*Δ, *ncs2*Δ*elp6*Δ*hel2*Δ and wild-type yeast. We observed pronounced ribosome collisions upstream of the strong stalling pairs CGA-AAA, CGA-CAA, GCA-AAA and GCA-CAA in *ncs2Δelp6*Δ cells (Fig. 4a and Supplementary Fig. 5b). Importantly, the extent of stalling was further increased in the *ncs2*Δ*elp6*Δ*hel2*Δ mutant (Fig. 4a and Supplementary Fig. 5b; red arrow), but not at sites of CGG-AAA or CGG-CAA (Supplementary Fig. 5c), confirming that RQC resolves ribosome collisions at strong stalling sites in *ncs2Δelp6*Δ cells.

**Fig. 4.**
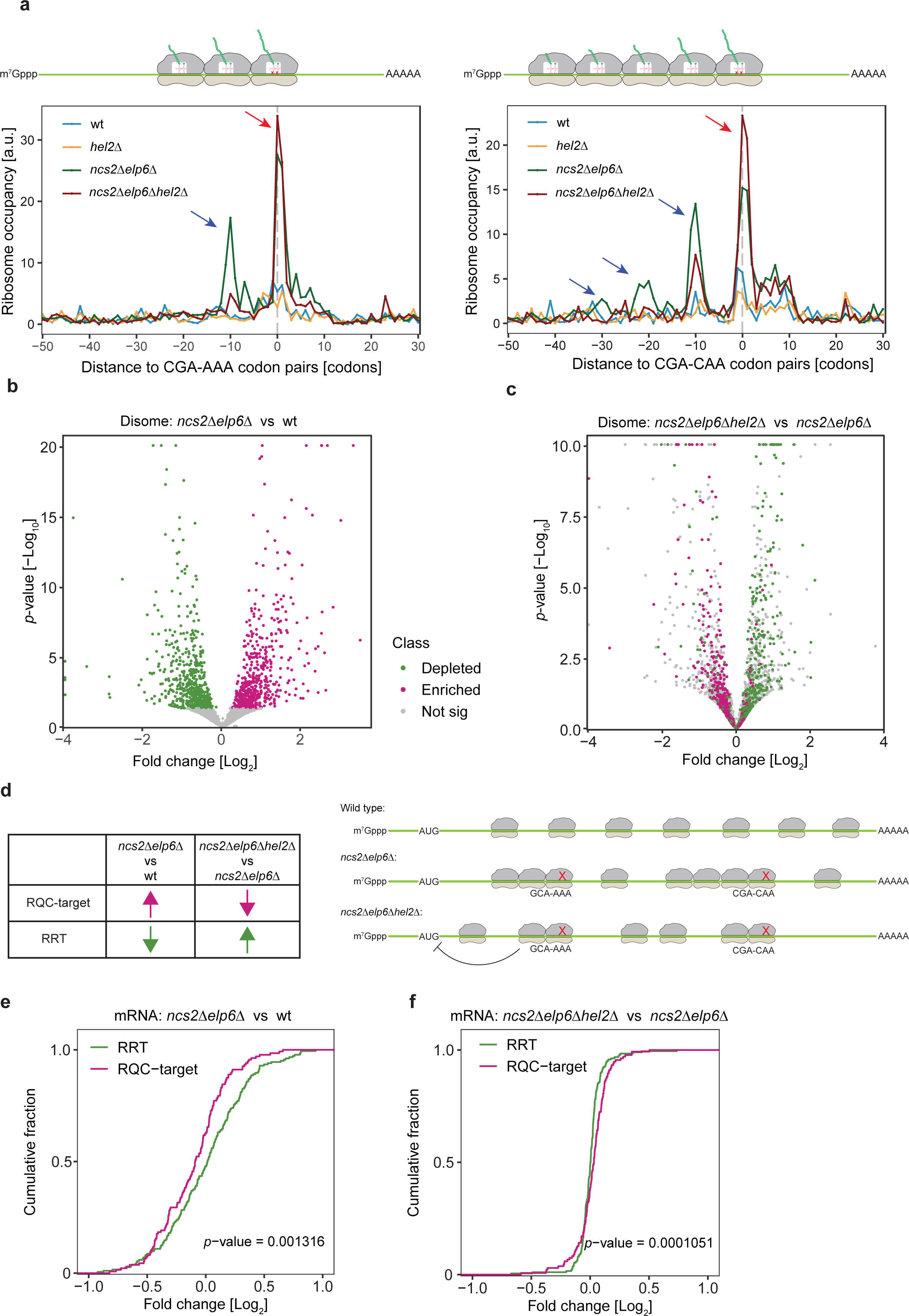
Ribosomes collide and are targeted by ribosome-associated quality control (RQC). (a) Disome occupancy around CGA-AAA (left) and CGA-CAA (right) codon pairs in wild-type, *hel2*Δ, *ncs2*Δ*elp6*Δ and *ncs2*Δ*elp6*Δ*hel2*Δ yeast. The average disome occupancy from two replicates was plotted. Peaks at position 0 represent the A site of the first stalling ribosome (red arrow). One queueing disome (blue arrow) was observed upstream of CGA-AA codon pairs at position -10 because the size of ribosome footprints is ∼10 codons (left). This indicates three ribosomes at the stalling site. Upstream of CGA-CAA codon pairs, three queueing disomes were observed at position -10, -20, and -30 in *ncs2*Δ*elp6*Δ, but only one queueing disome was found at position -10 in *ncs2*Δ*elp6*Δ*hel2*Δ. The cartoon above depicts collided ribosomes relative to the plot. (b) Differential expression of disome footprints comparing *ncs2Δelp6Δ* to wild type and comparing *ncs2*Δ*elp6*Δ*hel2*Δ to *ncs2*Δ*elp6*Δ using DESeq2^95^. Monosome footprints were used in the differential expression analysis for normalization due to a positive correlation between disome and monosome footprints (see Methods). Genes that show disome enrichment in *ncs2*Δ*elp6*Δ are highlighted in magenta while genes with less disomes are in highlighted in green. (c) Disome-enriched or -depleted genes in (b) are highlighted in the differential-expression analysis between *ncs2*Δ*elp6*Δ*hel2*Δ and *ncs2*Δ*elp6*Δ. (d) The strategy to assign genes as an RQC target (n=137) according to the disome enrichment in (b) and depletion in (c) or vice versa for RQC-refractory transcripts (RRTs) (n=188). Only genes significant (adjusted *p*-value < 0.05) for both criteria were selected. The schematic diagram on the right panel depicts the enrichment or depletion of ribosomes on mRNAs in three strains. (e) The change of mRNA levels comparing *ncs2*Δ*elp6*Δ to wild type was analyzed using DESeq2^95^. (f) Similar to (e) but comparing *ncs2*Δ*elp6*Δ*hel2*Δ to *ncs2*Δ*elp6*Δ. The Log_2_ fold change of RQC targets and RRTs was compared (one-sided Kolmogorov-Smirnov test).

Furthermore, we observed a strong signal of a third ribosome colliding with the disome for CGA-AAA, GCA-AAA, and even longer ribosomal queues for CGA-CAA (Fig. 4a and Supplementary Fig. 5b; blue arrow) in the *ncs2*Δ*elp6*Δ mutant. Interestingly, when *HEL2* is knocked out in *ncs2*Δ*elp6*Δ cells, ribosome queueing is reduced compared to the *ncs2*Δ*elp6*Δ mutant (Fig. 4a), indicating that ribosomal queueing is alleviated when the RQC pathway is inactivated. We posit that this effect is likely due to a downregulation of translation initiation, as we observed reduced rates of protein synthesis in *ncs2*Δ*elp6*Δ*hel2*Δ cells compared to the wild type and *ncs2*Δ*elp6*Δ mutants (Supplementary Fig. 5d). Similarly, and consistent with previous findings^35^, inactivation of RQC leads to activation of the ISR via Gcn2, resulting in increased translation of the transcription factor Gcn4 (Supplementary Fig. 5e) and transcriptional upregulation of its target genes (Supplementary Fig. 5f). In the absence of RQC, reduced translation initiation rates decrease the number of ribosomes on the transcript and reduce the likelihood of ribosome collisions.

Hel2-dependent ubiquitination marks ribosomes for splitting and subsequent degradation of the affected mRNA and nascent chains^34^. To identify specific transcripts that are targeted by the RQC pathway in U_34_-modification mutants, we compared disome enrichment between the wild type, *ncs2*Δ*elp6*Δ and *ncs2*Δ*elp6*Δ*hel2*Δ cells. First, we identified transcripts that show an increase in disomes in the *ncs2*Δ*elp6*Δ mutant compared to wild type, which indicates the occurrence of ribosome collisions (Fig. 4b). We found that these transcripts, enriched for disomes in the U_34_-modification mutant, were rescued in RQC-deficient *ncs2*Δ*elp6*Δ*hel2*Δ cells (Fig. 4c), consistent with reduced ribosomal queueing upstream of codon pairs in the *ncs2*Δ*elp6*Δ*hel2*Δ mutant (Fig. 4a). This analysis allowed us, for the first time, to identify endogenous RQC targets in a genome-wide manner. Therefore, we selected transcripts that are significantly upregulated in *ncs2*Δ*elp6*Δ yeast but significantly downregulated in *ncs2*Δ*elp6*Δ*hel2*Δ cells as RQC target transcripts, while we considered transcripts with the opposite pattern (downregulated in *ncs2*Δ*elp6*Δ but upregulated in *ncs2*Δ*elp6*Δ*hel2*Δ) as RQC-refractory transcripts (RRTs) and used them as a negative control (Fig. 4d). In total, we identified 188 RQC target transcripts and 137 RRTs (Supplementary Table 2). Using these two groups, we determined the frequency of modification-sensitive codons in each category and found that RQC targets are significantly enriched for non-optimal codons and codon pairs dependent on U_34_ modifications, confirming the accuracy of our classification (Supplementary Fig. 5g). However, whereas RTT are enriched for genes implicated in general metabolic processes such as amino acid biosynthesis or nucleoside metabolism (Supplementary Fig. 5h, left), RQC-targets are only weakly linked to functional categories in GO analyses (Supplementary Fig. 5h, right) suggesting that tRNA modifications are not used to functionally group transcripts in yeast.

mRNAs recognized by the RQC pathway are subsequently degraded by NGD^30,31^. Consistent with NGD-mediated degradation of RQC target transcripts, we observed that mRNA levels of RQC targets are significantly lower than RRTs in *ncs2*Δ*elp6*Δ cells when compared to wild type, whereas they are enriched in a 5PSeq dataset that we generated to quantify actively degrading mRNA (Fig. 4e and Supplementary Fig. 5i)^68^. Both effects are abolished in *ncs2*Δ*elp6*Δ*hel2*Δ cells (Fig. 4f and Supplementary Fig. 5i), demonstrating that mRNA levels of RQC targets are rescued in the absence of Hel2. Taken together, these findings show that U_34_-modification defects slow down ribosomes, trigger ribosome collisions, and activate RQC, which in turn degrades the affected mRNAs.

### The RQC pathway prevents U_34_-modification-dependent protein aggregation

A lack of U_34_ modifications triggers protein aggregation^9,29,69^. In light of our findings above, we asked how the observed ribosome collisions, RQC activation, and resultant mRNA degradation affect protein aggregation. Therefore, we reanalyzed a published inventory of proteins that aggregate in the absence of U_34_-modifications^9^. Of the 1080 proteins detected by mass spectrometry in *ncs2*Δ*elp6*Δ and wild-type yeast, we found that 610 proteins are more likely to aggregate in *ncs2*Δ*elp6Δ* cells compared to wild type, while 401 proteins are less likely to aggregate in the mutant strain, which we used as non-aggregating controls (See methods; Supplementary Table 3). We first asked whether aggregation-prone proteins are enriched in RQC targets or RRTs and found that aggregation-prone proteins are significantly enriched in RRTs compared to RQC targets, while we observed the opposite trend for non-aggregating proteins (Fig. 5a). This indicated that mRNA of non-aggregating proteins are more likely RQC target transcripts than mRNA of aggregation-prone proteins. Consistently, we detected less disomes in transcripts of aggregation-prone proteins when comparing *ncs2*Δ*elp6*Δ to wild-type cells, but elevated levels of disomes on the same transcripts in *ncs2*Δ*elp6*Δ*hel2*Δ cells lacking RQC activity (Supplementary Fig. 6a). Furthermore, we found that mRNA levels of non-aggregating proteins are reduced in *ncs2*Δ*elp6*Δ cells compared to wild type and that this effect is rescued in *ncs2*Δ*elp6*Δ*hel2*Δ cells (Supplementary Fig. 6b), indicating that RQC targets are less likely to form protein aggregates. Importantly, although we did not detect an enrichment of single U_34_-modification-dependent codons in either aggregation-prone or non-aggregating proteins (Supplementary Fig. 6c left), we found a significant enrichment of U_34_-modification-sensitive codon pairs in non-aggregating proteins (Supplementary Fig. 6c right). Since non-optimal codon pairs are more likely to cause ribosome collisions than single codons (Fig. 3d and Supplementary Fig. 4a and 4b), the enrichment of such codon pairs in non-aggregating proteins further supports our evidence that RQC targets are non-aggregating protein transcripts. Taken together, these findings suggest that the role of the RQC pathway extends beyond quality control of individual defective transcripts, but that it plays an additional role in preventing aggregate formation when translation dynamics are suboptimal. By degrading transcripts that accumulate collided ribosomes and potentially dysfunctional peptides, the RQC pathway reduces protein aggregation.

**Fig. 5.**
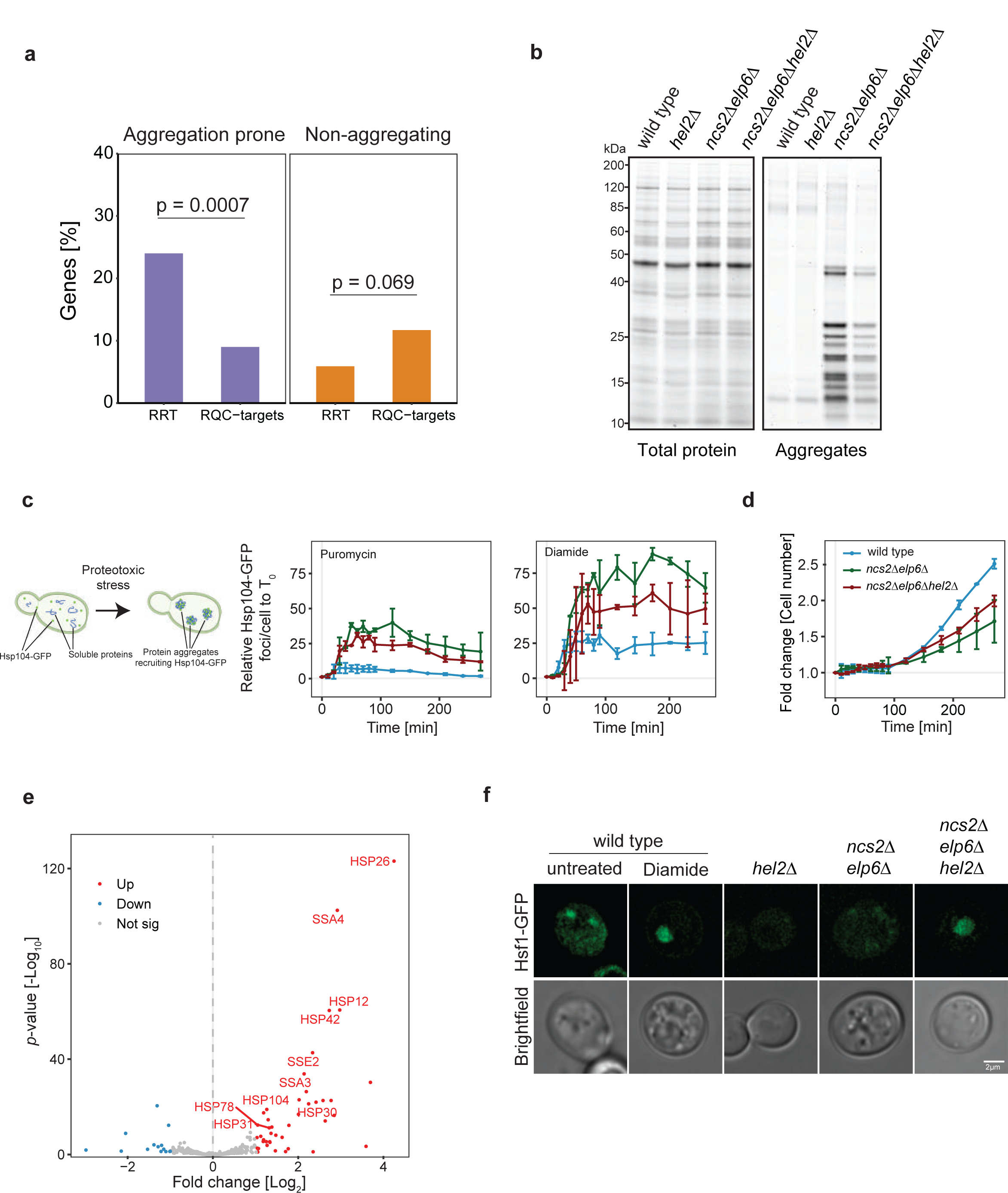
The RQC pathway alleviates protein homeostasis defects in coordination with chaperones. (a) Percentage of aggregation-prone (left) or non-aggregating (right) proteins in the RQC targets and RQC-refractory transcripts (RRTs) identified in Figure 5 (Fisher’s exact test). Proteins are considered as aggregation-prone in the mutant if enriched > 2-fold in the aggregates compared to wild type. (b) Total protein (left) and aggregates (right) in wild-type, *hel2*Δ, *ncs2*Δ*elp6*Δ and *ncs2*Δ*elp6*Δ*hel2*Δ yeast. (c) Yeast cells expressing Hsp104-GFP as readout for protein aggregation were exposed to 10 mM puromycin (left), 2 mM diamide (right) and the formation of GFP-positive foci was quantified over 4 hours. (d) Cell growth is monitored in Hsp104-GFP-based wild-type, *ncs2*Δ*elp6*Δ and *ncs2*Δ*elp6*Δ*hel2*Δ strains. (e) Differential expression analysis of mRNA levels comparing *ncs2*Δ*elp6*Δ*hel2*Δ to *ncs2*Δ*elp6*Δ. Upregulated genes with *p*-adjusted value < 0.05 and Log_2_ fold change > 1 are highlighted in red while downregulated genes with *p*-adjusted value < 0.05 and Log_2_ fold change < -1 are highlighted in blue. Significantly upregulated chaperones are labeled. (f) Hsf1-GFP was used to detect the localization of Hsf1 by live-cell microscopy (top row: GFP; bottom row: brightfield; scale bar for all images in this panel: 2 µm). As positive control, wild-type cells were treated with 2 mM diamide.

### The RQC Pathway is Coordinated with Molecular Chaperones

To understand the impact of RQC on protein homeostasis, we next asked how the absence of Hel2 affects protein aggregation in cells lacking U_34_ tRNA modifications and purified endogenous protein aggregates from wild-type, *hel2*Δ*, ncs2*Δ*elp6*Δ and *ncs2*Δ*elp6*Δ*hel2*Δ cells^9,28^. We expected to observe an increase of protein aggregates in *ncs2*Δ*elp6*Δ*hel2*Δ cells due to an accumulation of defective mRNAs and misfolded proteins. However, the levels of aggregating proteins were reduced in *ncs2*Δ*elp6*Δ*hel2*Δ cells compared to *ncs2*Δ*elp6*Δ yeast (Fig. 5b). To confirm this surprising finding, we performed high-content microscopy using Hsp104-GFP to monitor the formation of protein aggregates in response to chemically perturbed translation or protein stress *in vivo*. Similarly, we found that knocking out *HEL2* rescues the formation of Hsp104-GFP foci in *ncs2*Δ*elp6*Δ cells (Fig. 5c). Furthermore, removing Hel2 rescued the growth defect of U_34_-modification mutants (Fig. 5d), implying that additional quality-control systems contribute to the prevention of protein aggregation in the absence of functional RQC.

To identify the responsible pathways, we performed differential mRNA expression analysis between *ncs2*Δ*elp6*Δ*hel2*Δ and *ncs2*Δ*elp6*Δ yeast, which revealed a striking increase in the expression of molecular chaperones in *ncs2*Δ*elp6*Δ*hel2*Δ cells (Fig. 5e), and a gene ontology (GO) analysis for genes significantly upregulated in *ncs2*Δ*elp6*Δ*hel2*Δ cells similarly showed an enrichment of GO terms related to protein folding (Supplementary Fig. 6f). However, the expression of chaperones is not altered in *ncs2*Δ*elp6*Δ or *hel2*Δ cells compared to wild type (Supplementary Fig. 6d and 6e). This suggests that RQC monitors suboptimal translation dynamics antagonistically to molecular chaperones to maintain cellular protein homeostasis. The RQC pathway genetically interacts with Heat shock factor 1 (Hsf1), a key factor in maintaining protein homeostasis^37^. When protein homeostasis is normal, Hsf1 is sequestered in the cytoplasm by molecular chaperones. When protein misfolding occurs, chaperones bind to their client proteins and release Hsf1, which then translocates to the nucleus to upregulate chaperone expression^70^. Thus, the localization of Hsf1 can be used as a proxy for its chaperone-inducing activity. Therefore, we monitored the localization of Hsf1-GFP and found that it is mainly located in the cytoplasm in *hel2*Δ, *ncs2*Δ*elp6*Δ, and wild-type cells. However, it relocalizes to the nucleus in *ncs2*Δ*elp6*Δ*hel2*Δ cells (Fig. 5f). Furthermore, overexpression of Hsf1-GFP rescues the stress sensitivity of *ncs2*Δ*elp6*Δ cells (data now shown). These findings show that Hsf1 mediates the chaperone response in the absence of RQC, implying that the RQC pathways itself is typically enough to keep the heat-shock response at bay. Altogether, we conclude that the heat-shock response is a secondary cellular response that is activated when RQC fails to suppress translation-induced protein homeostasis defects.

## Discussion

Codon-specific translation rates, and therefore codon optimality, are generally determined by averaging ribosome occupancy of all instances of a codon across the transcriptome^8,9,71–74^. Here, we used high-resolution ribosome profiling to pinpoint translational slowdown events with high positional accuracy, revealing stark context-dependency to translational perturbations. Using yeast devoid of U_34_ tRNA modifications to probe the effects of codon-specific translational slowdown, we reveal a differential requirement for tRNA modifications between the ribosomal A site and P site, ribosome stalling at specific codon pairs that trigger ribosome collisions, and a hierarchical interplay between RQC and molecular chaperones to resolve translation impairments and protein homeostasis defects.

While previous low-resolution studies revealed slowdown during the decoding of AAA and CAA by their cognate mcm^5^s^2^U-modified 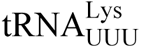 and 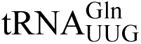, respectively in cells lacking U_34_ modifications^8,9^, our high-resolution analysis uncovered that ncm^5^U and mcm^5^U are required during wobble decoding of a subset of G-ending codons (AGG, CCG, GCG, GGG, and GUG; Fig. 1d and Supplementary Fig. 1c) in the ribosomal A site. Unexpectedly, we also find that ncm^5^U is required for efficient processing of cognate codons (ACA, GCA, GUA, UCA) in the P site (Fig. 2c and 2d and Supplementary Fig. 3c and 3d). This demonstrates that the same modification can differentially tune the anticodon-codon interaction of different codons in the A and P site of the ribosome. To our knowledge, such an effect has not been reported previously.

Importantly, we observed that certain instances of codons are more vulnerable to modification defects compared to other codons of the same type - even within the same transcript. We systematically evaluated factors that might explain the observed vulnerability and found that this phenomenon largely depends on the P-site codon that synergistically deoptimizes a suboptimal A-site codon (Fig. 2). Codon pair biases have been found in many organisms, where suboptimal codon pairs can compromise translation rates^36,75,76^ or contribute to pausing, reduced gene expression and decreased mRNA stability^10,77^. When analyzing slow codon pairs in U_34_-modification mutants, we found that U_34_-related codons and CGA in the P site enhance the slowdown of A-site codons. CGA codons are commonly observed in slow pairs, which has been explained by a change in mRNA conformation in the P site^10,57^. We hypothesize that the absence of ncm^5^U or mcm^5^U during A-U pairing may similarly affect mRNA structure at the P site and reduce the binding efficiency of hypomodified tRNA at the A site. Furthermore, our data argue that the phenotypes of tRNA modification mutants are likely mediated by codon-pair defects. The identification of codon pairs that become suboptimal under specific conditions is critical, as these cannot be predicted from their frequency in the transcriptome and may differ when tRNA modification levels are altered in response to environmental conditions. However, we found no evidence that such suboptimal codon pairs are used by cells to tune translation dynamics in a modification-dependent manner.

We show that ribosomes stall at suboptimal codon pairs leading to ribosome collisions (Fig. 2, 3b, 4a and Supplementary Fig. 3c, 3d, 4a and 5b), thereby triggering RQC as indicated by Hel2-dependent ubiquitination of ribosomal proteins (Supplementary Fig. 5a). Depletion of Hel2 resulted in increased disome occupancy at modification-dependent codon pairs in the absence of U_34_ modifications, indicating that Hel2 resolves collided ribosomes at such stalling sites. Interestingly, in *ncs2*Δ*elp6*Δ*hel2*Δ cells, we observed less ribosome queueing upstream of CGA-CAA and CGA-AAA pairs and fewer disome footprints at affected transcripts (Fig. 4a, 4b and Supplementary Fig. 5b). This counterintuitive finding suggests that the number of colliding ribosomes is much higher than the number of stalling ribosomes that cause these collisions. At the same time, translation initiation is inhibited to reduce the ribosome load on mRNA, which alleviates ribosomal queueing. Persistent ribosome collisions activate the *GCN2*-mediated stress response pathway, leading to a block in translation initiation^78^. Consistent with this, we observed reduced initiation in *ncs2*Δ*elp6*Δ*hel2*Δ cells and a striking upregulation of Gcn4 targets (Fig. 4a and Supplementary Fig. 5b and 5e), indicative of Gcn2 activation. Also in line with our results, it has been shown that RQC and ISR are antagonistic pathways that compete for collided ribosomes as their substrate^39,79^. Therefore, deletion of *HEL2* promotes increased Gcn2 activation^35^.

The combination of tRNA modification defects and RQC impairment induced a striking upregulation of molecular chaperones that we did not observe when either pathway was perturbed independently (Fig. 5e). This shows that the different quality control systems work hierarchically to ensure the synthesis of correctly folded proteins. Our study reveals that ribosomal stalling at slow codon pairs triggers the RQC pathway and that the nascent peptides of RQC target genes are likely degraded (Fig. 6). In addition to monitoring mRNA quality, we show a novel role for RQC in preventing general protein aggregation by targeting suboptimal translation events at an early stage. If stalled ribosomes cannot be resolved by RQC, the defective nascent peptides have a high potential to interact with other proteins and form aggregates^80^. Importantly, the fact that chaperones are only slightly upregulated in *ncs2*Δ*elp6*Δ yeast suggests that the Hsf1-mediated chaperone response is suppressed when RQC is active^37^. RQC and HSR likely act antagonistically for economic reasons since chaperones require high levels of ATP. Under physiological conditions, RQC efficiently removes potentially misfolding proteins by simply degrading mRNA and the nascent chain, allowing subsequent recycling of the different components of the translation machinery. The role of RQC to recognize defects in translation dynamics and to avoid co-translational misfolding may become even more critical during the co-translational assembly of protein complexes by subsequent ribosomes^81,82^. Although we have used yeast as a model system in this study, similar mechanisms are likely to occur in metazoans. For example, mutations of several tRNA modifying enzymes but also the RQC machinery are associated with neurodegenerative phenotypes^20,21,83^. Since neurons harbor complex proteostasis needs, coordination between quality-control pathways may contribute to the integrity of brain cells and prevention of neurodegeneration.

**Fig. 6.**
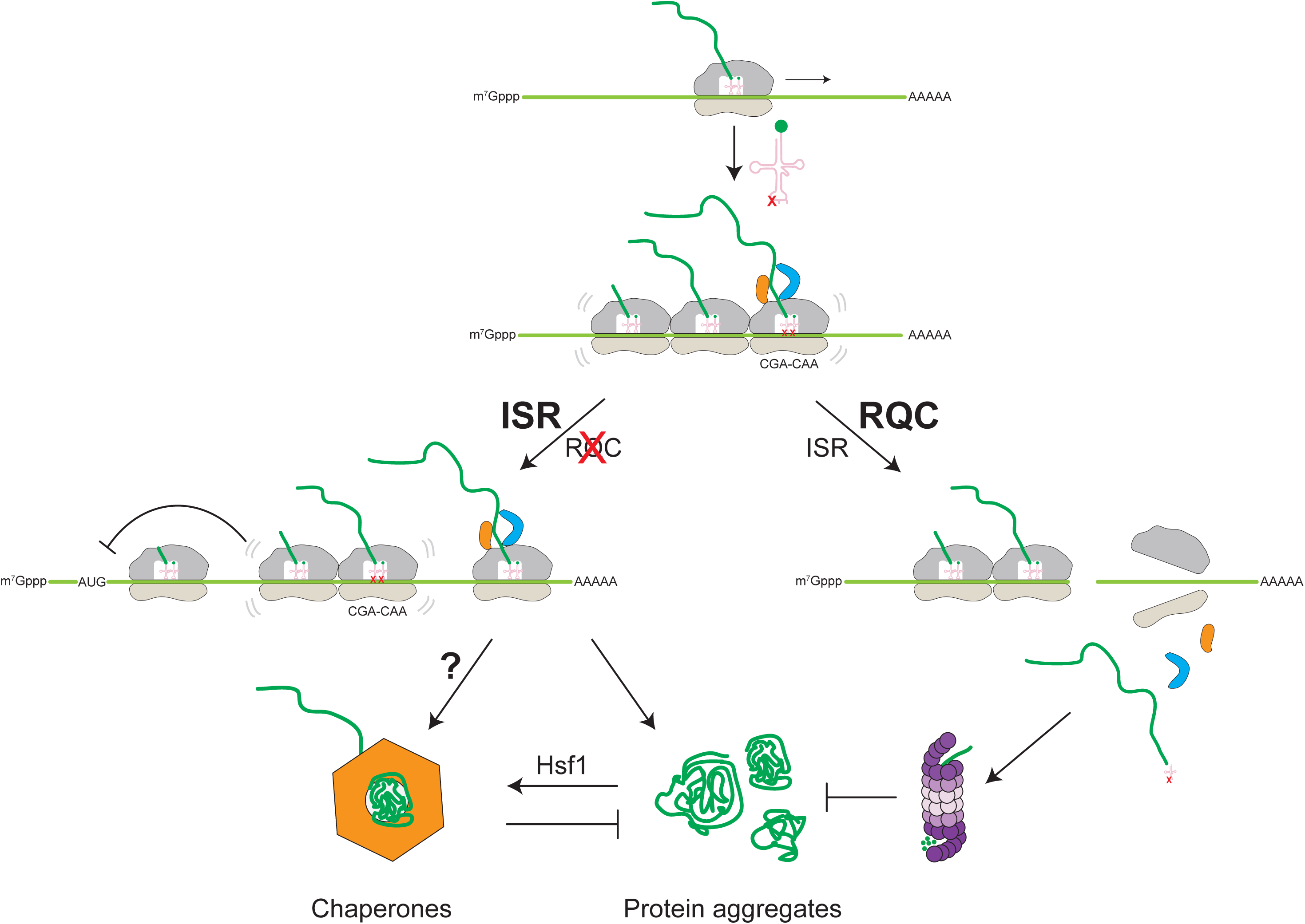
Working model. Model for the cooperative surveillance of the cellular quality-control systems in response to codon-specific translation defects induced by hypomodification of tRNA. When tRNA modifications are absent, ribosomes collide at modification-dependent codon pairs. This induces the ribosome-associated quality control (RQC) pathway, which targets the aberrant transcript (right). Peptides synthesized by stalling ribosomes are degraded by the proteosome and ribosome subunits recycled. Thus, RQC targets are less likely to form protein aggregates. When the RQC is inactive (left), the integrated stress response (ISR) is strongly induced and downregulates translation initiation to prevent further collisions and the cell overexpresses chaperones via Hsf1 to reduce protein aggregation.

In conclusion, we show that U_34_ tRNA modifications differentially affect A-site and P-site codons, highlighting the importance of codon pairs for context-dependent codon optimality. Seemingly mild codon-specific translation defects can induce ribosome stalling and collisions through the synergistic effect of suboptimal codons. While RQC is able to correct mild translation defects, it is overwhelmed when such defects become persistent. When translation dynamics are perturbed, ribosomes deviate from the optimal balance between protein synthesis rates and cotranslational folding, forcing cells to remedy this perturbation by integrating multiple branches of cellular quality control. Although our study focused on U_34_ modifications, this methodology and conceptual framework can be extended to other factors that alter codon optimality, such as mutations in the translation machinery, tRNA copy number variations, or epitranscriptomic modifications. Therefore, the analysis of suboptimal codon pairs will likely be key for the design of expression constructs, including RNA therapeutics.

## Material and Methods

### Yeast strains and growth conditions

Experiments were performed in S288c (BY4741) yeast. All yeast strains and plasmids are listed in Supplementary Tables 4 and 5. Yeast cultures were grown to mid-exponential phase OD_600_ 0.4–0.5 at 30 °C, 200 rpm in YPD (Formedium).

### Ribosome Profiling

Ribosome profiling was performed as described with minor modifications^43,44^. Yeast cells were harvested by vacuum filtration and the pellet snap frozen in liquid nitrogen with droplets of lysis buffer. The mixture of cells and buffer was lysed using a freezer mill (Spex SamplePrep 6770). The resulting powder was thawed in a water bath and cleared by 2 rounds of centrifugation (3’000 g for 3 min at 4 °C and 10’000 g for 5 min at 4 °C). A_260_ absorbance of the lysate was measured, and 8 U of RNA were digested with 800 U RNaseI (AM2294, Invitrogen) in 80 µL reaction volume at 22 °C and 1’400 rpm for 1 h.

Digested samples were loaded on a 10 %-50 % sucrose gradient and separated by ultracentrifugation at 35’000 rpm, 4 °C for 3 h using a SW41 rotor (Beckman Coulter). Sucrose gradients were fractionated using a Piston Gradient Fractionator (Biocomp). Both monosome and disome fractions were collected. SDS was added to a final concentration of 1 % prior to snap freezing the fractions. RNA was extracted from the fractions using hot acid phenol-chloroform. 10 µg of RNA per sample was size-separated on a 15 % TBE-Urea polyacrylamide gel. For monosomes, the gel between the 28 nt and 32 nt marker was excised and for disomes between 56 nt and 64 nt. The excised bands were crushed in a 1.5 mL microtube with 400 µL elution buffer (0.3 M NaOAc pH 5.5, 1 mM EDTA pH 8.0, 0.1 U/µl SuperaseIN (AM2694, Invitrogen) and eluted overnight at 4 °C on a spinning wheel. The next day, the gel pieces were removed using a Spin-X column (CLS8162, Merck, Corning Costar Spin-X centrifuge tube filters) and RNA was precipitated overnight at -80 °C.

Ribosome-protected fragments were dephosphorylated using T4 Polynucleotide Kinase (M0201, New England Biolabs). Dephosphorylated RNA was extracted using ROTI Acqua P/C/I for RNA extraction (X985.1, Carl Roth, ROTI Aqua-P/C/I) and RNA precipitated with NaOAc overnight at -80 °C. Subsequently, RNA was ligated using T4 RNA ligase 2 (M0242, New England Biolabs) to a preadenylated linker and purified with ROTI Aqua P/C/I for RNA extraction and precipitated with NaOAc. Samples were reverse transcribed using SuperScript III (18080093, Invitrogen). Subsequently, template RNA was hydrolyzed using NaOH. cDNA was run on a 10 % TBE-Urea polyacrylamide gel. The band corresponding to the cDNA was excised and crushed in 360 µL ddH_2_O and eluted for 20 min in a thermoblock at 70 °C and 1’400 rpm. Gel pieces were removed using a Spin-X column and the cDNA precipitated with NaOAc overnight at -80 °C and resuspended in 11 µL ddH_2_O. cDNA was circularized with CircLigase II (CL9021, Lucigen). Libraries were PCR amplified using Phusion DNA Polymerase (M0530, New England Biolabs). Quality assessment of the libraries and sequencing were performed by the NGS platform of the University of Bern.

### mRNA and 5P Sequencing

Total RNA extraction was performed as described in^9^. A fraction of the cells harvested for Ribosome profiling was used for mRNAseq. The frozen pellets were resuspended in lysis buffer (50 mM NaOAc pH 5.5, 10 mM EDTA, 1 % SDS) and cells were lysed using a Fastprep24 bead beater. Lysates were clarified by centrifugation. Samples were treated with 100 µg/mL Proteinase K (AM2546, Invitrogen) for 20 min at 60 °C. Finally, RNA was extracted with acid-phenol/chloroform. Library preparation was performed using a TruSeq-Stranded-mRNA kit (20020594, Illumina) according to the manufacturer’s instructions. 5Pseq libraries were generated as described in^84^.

### Fluorescence Measurement of the Reporter

2 to 3 OD_600_ of yeast cultures of OD_600_ 0.6-0.8 were pelleted and resuspended in lysis buffer (20 mM MES, 100 mM NaCl, 30 mM Tris-HCl pH 7,5, 1 % Triton X-100, supplemented with 1x cOmplete protease inhibitor (4693159001, Merck). Cells were lysed using a Fastprep24 bead beater and the lysate was cleared by centrifugation. GFP and mKate fluorescence was measured using an Infinite M1000 plate reader (Tecan).

### ^35^S labeling

3 OD_600_ units of yeast during exponential growth were inoculated to 3 ml SC -Met medium containing Glucose and 15 μCi/ml ^35^S L-methionine. 500 µl aliquots were taken at timepoints 0 min, 10 min and 30 min. Cells were pelleted and resuspended in 500 ml ice-cold 20 % TCA, incubated on ice for 20 min, precipitated and washed with acetone. Samples were run on a 4-12 % gradient gel (mPAGE, SigmaAldrich) and radioactive signals were scanned using the TyphoonFLA1000 Phosphoimager. Bands were quantified using AzureSpot Analysis Software (Azure Biosystems).

### Live-cell imaging

Yeast cells were grown in log-phase in a high-cell-density medium (HCD: MSG 4.5 mg/ml, YNB 12 mg/ml, Inositol 1.8 µg/ml, 20 mM MES buffer pH 6.0, 6 % glucose and amino acids)^85^ at 30 °C. Overnight cultures were diluted to an OD_600_ of 0.3 and further incubated until OD_600_ = 0.6. To monitor the translocation of Hsf1-GFP to the nucleus, cultures were treated with 2 mM diamide for 1 h. For nuclear staining, the samples were incubated with 100 nM Hoechst reagent (H-1399, Invitrogen) for 5 min. Cells were then pelleted at 3’000 g for 3 min and resuspended in fresh HCD medium. Yeast cells were attached to 35 mm glass bottom dishes (D35-20, 1.5, In Vitro Scientific) pretreated with concanavalin A type IV (1 mg/ml, Sigma-Aldrich), and imaged at room temperature. Fluorescent microscopy images were recorded with a DeltaVision Ultra High Resolution Microscope with UPlanSApo 100 ×/1.4 oil Olympus objective, using a sCMOS pro.edge camera (GE Healthcare, Applied Precision). Image analysis was performed using FIJI^86^. Images from each figure panel were taken with the same imaging setup and are shown with identical contrast settings. All images were generated by collecting a z-stack of 25 pictures with focal planes 0.20 μm apart. Single focal planes of representative images are shown.

### High-content microscopy (Hsp104 assay)

Yeast cells expressing *HSP104-GFP* from the endogenous locus were grown in SD-complete medium until reaching OD_600_ ∼ 0.6. The cells were then diluted to an OD_600_ = 0.1 and 20 µl of the diluted culture were seeded to each well of a 384-well plate with a transparent bottom.

Cell suspensions were centrifuged at 2’000 g for 1 min to form a monolayer. Subsequently the plate was imaged on an Operetta High Content Analysis System (PerkinElmer) for 4 h. Images were captured every 10 min for the first 90 min and subsequently every 30 min. Two different fields of view (3 planes each) were captured with digital phase contrast and GFP channels for each well and time point using a 60x high NA air objective. The images were quantified using Harmony Software v4.1, Revision 128972 (PerkinElmer). Briefly, all planes were stacked, and cells were filtered based on their roundness and texture using the phase-contrast images, as well as GFP background intensity. For quantification of GFP-positive spots, size, intensity, and relative contrast to the surrounding area were taken into account. For each condition, between 500 and 1000 cells were imaged at time point T0.

### Cryo-EM single-particle reconstruction

Cryo-EM datasets were collected at the National Cryo-EM Centre SOLARIS (Kraków, Poland). The datasets contained 8595 (WT#1), 10009 (WT#2), 9486 (*ncs2*Δ*elp6*Δ#1) and 8583 (*ncs2*Δ*elp6*Δ#2) movies. The movies were acquired using a Titan Krios G3i microscope (Thermo Fisher Scientific) operated at 300 kV accelerating voltage, magnification of 105k, and corresponding pixel size of 0.8456 Å/px. A K3 direct electron detector used for data collection was fitted with BioQuantum Imaging Filter (Gatan) using a 20 eV slit. The K3 detector was operated in counting mode. Imaged areas were exposed to 40 e^−^/Å^2^ total dose (corresponding to ∼16 e^−^/px/s dose rate measured in vacuum). 40-frame movie stacks were obtained using under-focus optical conditions with a defocus range of −2.2 to −0.7 µm and 0.3 µm steps. The collected datasets were analyzed using cryoSPARC v4.4.1^87^. First, patch motion correction and patch CTF estimation steps were performed. Next, a template picking based on a consensus *S. cerevisiae* 80S ribosome volume, and subsequent 2D classification, resulted in 858868 (WT#1), 758909 (WT#2), 976463 (*ncs2*Δ*elp6*Δ#1) and 672666 (*ncs2*Δ*elp6*Δ#2) particles, respectively. Following "Ab-initio Reconstruction", and "Homogenous Refinement", the particles served as an input for 3D classification. To optimize class distribution for comparative analysis, the particles from 4 datasets (WT#1 + WT#2 + *ncs2*Δ*elp6*Δ*#*1 + *ncs2*Δ*elp6*Δ*#*2) were mixed into a common pool. First, "Heterogenous Refinement" was used on the common pool to remove particles corresponding to 60S complexes. Next, to focus the analysis on 80S translating ribosomes in different states, focused 3D classifications were performed using soft masks generated from regions occupied by E-, P-, and A-site tRNA, and EF2 (Supplementary Fig. 2c). After each class selection step, the particles originating from individual strains and replicates were enumerated using the intersection option of the Particle Sets tool. The final particle stacks were reextracted without binning for "Reference Based Motion Correction" in cryo-SPARC^88^. During final rounds of "Homogeneous Refinement", the particles were subjected to "Defocus Refinement", "Global CTF Refinement", and "Edwald Sphere Correction" to generate high-resolution maps. Local map resolution was calculated using cryoSPARC (Supplementary Fig. 2d and 2e). The combined half-maps and corresponding refinement masks were used for DeepEMhancer sharpening^89^. The cryo-EM maps were displayed using ChimeraX version 1.7^90^.

### Protein aggregate purification

Protein aggregates were isolated as described in^28^. 50 O.D._600_ of exponentially growing cells were harvested by vacuum filter and immediately snap-frozen. The pellets were resuspended in 1 ml of lysis buffer (20 mM NaPi pH=6.8, 1 mM EDTA, 10 mM DTT, 0.1 % Tween 20) supplemented with protease inhibitors (0.5 mM AEBSF, 10 µg/ml aprotinin, 0.5 mg/ml benzamidine, 20 µM leupeptin, 5 µM pepstatin A). 60 U of zymolyase T20 (Zymo Research) were added, and cells were incubated at 22 °C for 20 min at 1’400 rpm. Extracts were chilled on ice and sonicated eight times with 50 % amplitude and duty cycle 50 % with a FisherBrand 120 tip sonicator (Thermo Fisher). Cell debris was pelleted by centrifugation at 200 g, 4°C for 20 min. Protein concentration of the supernatants was quantified using the Bio-Rad Protein Assay. To precipitate aggregates, 2 mg of total protein in 1 ml of lysis buffer were centrifuged at 16’000 g, 4°C for 20 min. The supernatant was discard and the pellet resuspended in washing buffer I (20 mM NaPi pH=6.8, 2% NP-40, protease inhibitor cocktail) and sonicated six times with amplitude 50 %, duty cycle 50 %, and precipitated again at 16’000 g, 4 °C for 20 min. The pellet was again resuspended in washing buffer I, sonicated and precipitated as before. Finally, aggregates were resuspended in detergent-free washing buffer by sonicating 4 times with amplitude 35 %, duty cycle 50 %. Following precipitation, pellets were dissolved in Laemmli sample buffer with 100 mM DTT and 8M urea and boiled at 95 °C for 5 min. Total protein extracts and aggregates were run on 4-12 % NuPAGE Bis-Tris gels (Life Technologies) in MES-SDS running buffer and gels were stained with AzureRed Fluorescent Protein Stain (Azure Biosystems). Gels were scanned using a Sapphire Biomolucar Imager (Azure Biosystems) and images quantified with AzureSpot (Azure Biosystems).

### Ribosome Mass Spectrometry

50 OD_600_ units of yeast during exponential growth were harvested by vacuum filtering and snap-frozen with droplets of lysis buffer^44^. The mixture of cells and buffer was lysed using a freezer mill (Spex SamplePrep 6770). The resulting powder was thawed in a water bath and cleared by 2 rounds of centrifugation. A_260_ absorbance of the lysate was measured, and 15 U of RNA were diluted 1:1 with 2 mM CaCl_2_ and digested with 100 U Micrococcal Nuclease (Thermo Scientific™ EN0181) at 22 °C and 1’400 rpm for 15 min. Digested samples were loaded on a 10-50 % sucrose gradient and separated by ultracentrifugation at 35’000 rpm, 4 °C for 3 h using a SW41 rotor (Beckman Coulter). Sucrose gradients were fractionated using a Piston Gradient Fractionator (Biocomp). The disome peak was collected and proteins precipitated using TCA.

### Sequencing Data Processing

Sequencing data from ribosome profiling was pre-processed by removing the adapter using the Fasx_toolkit. Four randomized linker nucleotides at the 3’ end were trimmed. If the library was generated by dual randomization, three nucleotides at the 5’ end and six at the 3’ end were trimmed. The quality of data was evaluated using FastQC (v0.11.7). Processed reads were mapped to non-coding RNA, including rRNAs, tRNAs, snoRNAs and other annotated lncRNAs using STAR (v2.5.3a) to remove non-protein-coding reads^91^. Remaining reads were subsequently mapped to all annotated open reading frames in yeast using bowtie (v1.2.3) with unique mapping mode allowing at most one nucleotide mismatch with parameters "-v 1 -m 1 --norc --best --strata"^92^. Each ORF was extended 21 bp at both 5’ end and 3’ end for the alignment of footprints around the start codons and stop codons. If an extra mismatched T was found at the 5’ end of the read, which is due to the extra insertion of an A by reverse transcriptase during the RT step, it was also removed. The processed data were used for downstream analysis.

The meta-analysis around the start codons or stop codons were performed by aligning the A of all AUG start codons at the 0 position or aligning the last nucleotide of all stop codons (UG<u>A</u>/UA<u>A</u>/UA<u>G</u>) at -1 position, respectively. Footprints in frames 0, 1 and 2 were defined, if the 5’ end of the read was mapped to the first, second and third nucleotide of the codon, respectively. Footprints are grouped by read length to present the 3-nt periodicity. Due to the incomplete or excessive digest of RNaseI, the assignment of the A site for each read length is determined according to the first high peak at around -12 nt. In our study, we selected 28 nt footprints with frame 0 (A-site offset: 16 nt) and 29 nt footprints with frame 0 (A-site offset 16 nt) and frame 2 (A-site offset: 17 nt) for downstream analyses of codon translation rates.

Disome profiling data were pre-processed like monosome profiling. 56-64-nt-long footprints are kept for analysis. Since no disome enrichment can be found around the start codon, when selecting the reads for downstream analysis, offsets were chosen according to the last disomes at the stop codons. RNA-seq reads were mapped to CDS without adaptor trimming.

### Calculation of Translation Rate of Codons

1. Read-based (Fig. 1d): The frequency of codons in the ribosomal A sites were quantified according to the length-specific offsets for all selected reads. The translation rate of sense 61 codons comparing mutant to wild type was calculated as^9^ by normalizing the frequency of codons in the A sites to that in the +1, +2, +3 sites, followed by calculating the ratio of translation rate between mutant and wild type (15 codons were excluded at both end of CDS). To avoid that few extremely affected codons shift other codons away from the mean (log_2_ ratio=0), the ratio of each codon is further normalized to the median of all ratios.
2. Position-based^93^ (Supplementary Fig. 1c): After assigning footprints to the A-site codon according to the selected offset, the read counts for each codon in the CDS can be quantified (15 codons at both 5’ and 3’ end of CDS were excluded). Only genes with read coverage of 0.1 reads per codon are included in the analysis. The pause score for each codon in each CDS is subsequently calculated by normalizing the read count of this codon to the average read count of all codons in this CDS:

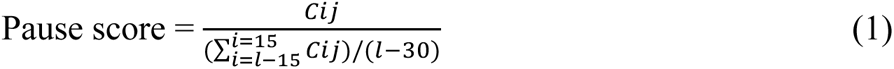

Where Cij is the read count of codon i in gene j, l is the length of CDS (codons). Vulnerability score of each codon in the mutant compared to wild type is calculated by normalizing the mutant pause score to that in wild type:

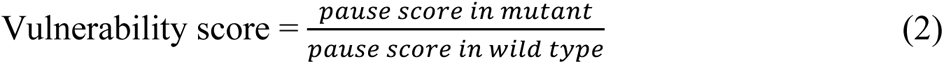

To avoid dividing by zero during the calculation of the vulnerability score, we added 1 read to each position before calculating the pause score in the genes that were selected for analysis (0.1 reads per codon). The median of vulnerability scores of three replicates in mutant and wild type is used to represent the susceptibility of this codon to the absence of U_34_ modifications.

### Motif Analysis

Vulnerability scores of each of the 61 sense codons are ranked and the top 1000 of the codons with high vulnerability scores were selected for motif analyses. Motif analysis was performed for the 30 codons upstream of the codons of interest using kpLogo and probability logo is shown^94^. Positions with significant enriched or depleted residues are highlighted in red (Bonferroni-corrected *p*-value < 0.01). To remove potentially embedded biases, the same analysis pipeline was implemented comparing two wild-type replicates. Upstream sequences of top 1000 codons were selected as the input of background sequence list in kpLogo.

### Di-codon Analysis

The pause score for each codon pair in each ORF was calculated as in formula (1). The average of pause scores of each codon pair in the ORFs was used as translation rate of this codon pair. The ratio of averaged translation rates from three replicates between mutant and wild type were calculated as the alteration of the translation rate of codon pairs. To visualize the change of all codon pairs, codon pairs with ratio > 2 are all assigned to 2 in the di-codon plot. In the di-codon plot, the size of circle represents the number of codon pairs in all CDSs in the analysis.

### Analysis of Translation Near Codons of Interest

All specific codon/codon pairs are aligned at position 0. The pause scores upstream or downstream of the codons/codon pairs of interest were averaged with all codon/codon pairs within that position^93^. Disome footprints are assigned to the position according to the location of the A site of the stalling ribosome (45 nt downstream of the 5’ end of the disome read).

### Analysis of Disome Enrichment

The monosome/disome footprints uniquely mapped to each ORF were quantified using a custom script. The differential expression analysis of disome enrichment was performed using DESeq2 (v1.28.1)^95^. Since the disome profile is positively correlated with the monosome profile, the expression quantified by monosome footprints are included in the DE analysis using design = ∼ sampleType+condition+condition:sampleType. The likelihood-ratio test was used to test a reduced model (∼ sampleType+condition) and the significant genes with disome enrichment or depletion were filtered using adjusted *p*-value < 0.05. Genes with significantly enriched disomes in *ncs2*Δ*elp6*Δ but significantly decreased disomes in *ncs2*Δ*elp6*Δ*hel2*Δ were selected as RQC targets. Genes with significantly depleted disomes in *ncs2*Δ*elp6*Δ but significantly enriched disomes in *ncs2*Δ*elp6*Δ*hel2*Δ were selected as RQC-refractory transcripts (RRTs).

### DE Analysis of RNA-seq Data

The change of mRNA levels between strains was analyzed using DESeq2^95^. Significant genes were tested using a Wald test and filtered with adjusted *p*-value < 0.05. When performing differential-expression analysis between *ncs2*Δ*elp6*Δ*hel2*Δ and *ncs2*Δ*elp6*Δ mutants, we sought to correct for changes in transcription that may be caused by the depletion of Hel2. Hence, we included RNA-seq of *hel2*Δ and wild-type strains in the analysis as in the analysis of disome enrichment. However, the results were not altered whether including *hel2*Δ in the analysis or not.

### Analysis of 5PSeq Data

Adapter sequences were trimmed by cutadapt using parameters "-m 10 -a GATCGGAAGAGCACACGTCTGAACTCCAGTC"^96^. Untrimmed reads were also retained for downstream analysis due to the long insert size. The processed reads were mapped to non-coding RNA using STAR (v2.5.3a)^91^. The remaining reads were mapped to all annotated open reading frames in yeast using bowtie (v1.2.3) using the same parameters as for the analysis of ribosome profiling data^92^. PCR duplicates were further removed using UMI-tools with 8 nt randomized barcodes at the 5’ end of sequencing reads^97^. The 3-nt periodicity of reads was also evaluated. Unique mapped reads were quantified using a custom script. DE analysis was performed with DEseq2 (v1.28.1) by combining RNA-seq data using design = ∼ sampleType+condition+condition:sampleType^95^. The likelihood-ratio test was used to test a reduced model (∼ sampleType+condition). Log_2_ fold change of RQC-target and RRTs were compared between different strains.

### Additional Computational Analyses

The gene-ontology analysis was performed using the online Panther server on the gene-ontology resource (http://geneontology.org/). Genes for aggregate-related analysis are retrieved from^9^. Genes with fold change > 2 between *ncs2*Δ*elp6*Δ and wild type in aggregate proteomics are selected as aggregation-prone proteins and fold change < 0.5 as non-aggregating proteins.

## Acknowledgements

We thank Claudia Gräf, Michal Rawski, Paulina Indyka, Grzegorz Wazny and Marcin Jaciuk for support; Eric Westhof and all members of the Leidel lab for critical discussions and comments on the manuscript. We would like to thank the Next Generation Sequencing Platform for an excellent service in performing the high-throughput sequencing experiments and the Core Facility Proteomics & Mass Spectrometry of the University of Bern for performing our proteomics experiments. This work was supported by the Canton of Bern and grants by the Swiss National Science Foundation [310030_184947 to S.A.L.]; the German Research Foundation [Project ID 409673687], SFB 1381 [Project ID 403222702], SFB 1177 [Project ID 259130777], Germany’s Excellence Strategy [CIBSS - EXC-2189 - Project ID 390939984]; the European Research Council (ERC) under the European Union’s Horizon 2020 research and innovation programme [Project ID 769065 to C.K., Project ID 669168 to H.R.S., and Project ID 101001394 to S.G]. O.S. was supported by a Marie Sklodowska-Curie Postdoctoral Fellowship (European Commission) [Grant ID 746340] and a Rubicon Postdoctoral Fellowship (Netherlands Organisation for Scientific Research) [Grant ID 019.162LW.028]. This publication was partially developed under the provision of the Polish Ministry and Higher Education project "Support for research and development with the use of research infrastructure of the National Synchrotron Radiation Centre SOLARIS" under contract nr 1/SOL/2021/2. We gratefully acknowledge the Polish high-performance computing infrastructure PLGrid (HPC Centers: ACK Cyfronet AGH) for providing computer facilities and support within computational grant no. PLG/2020/014009.

This work reflects only the authors’ view, and the funding organizations are not responsible for any use that may be made of the information it contains.

The authors declare no competing financial interests.

