## Supplementary Figures for "tRNA modifications tune decoding of codon pairs to prevent cellular quality control responses"

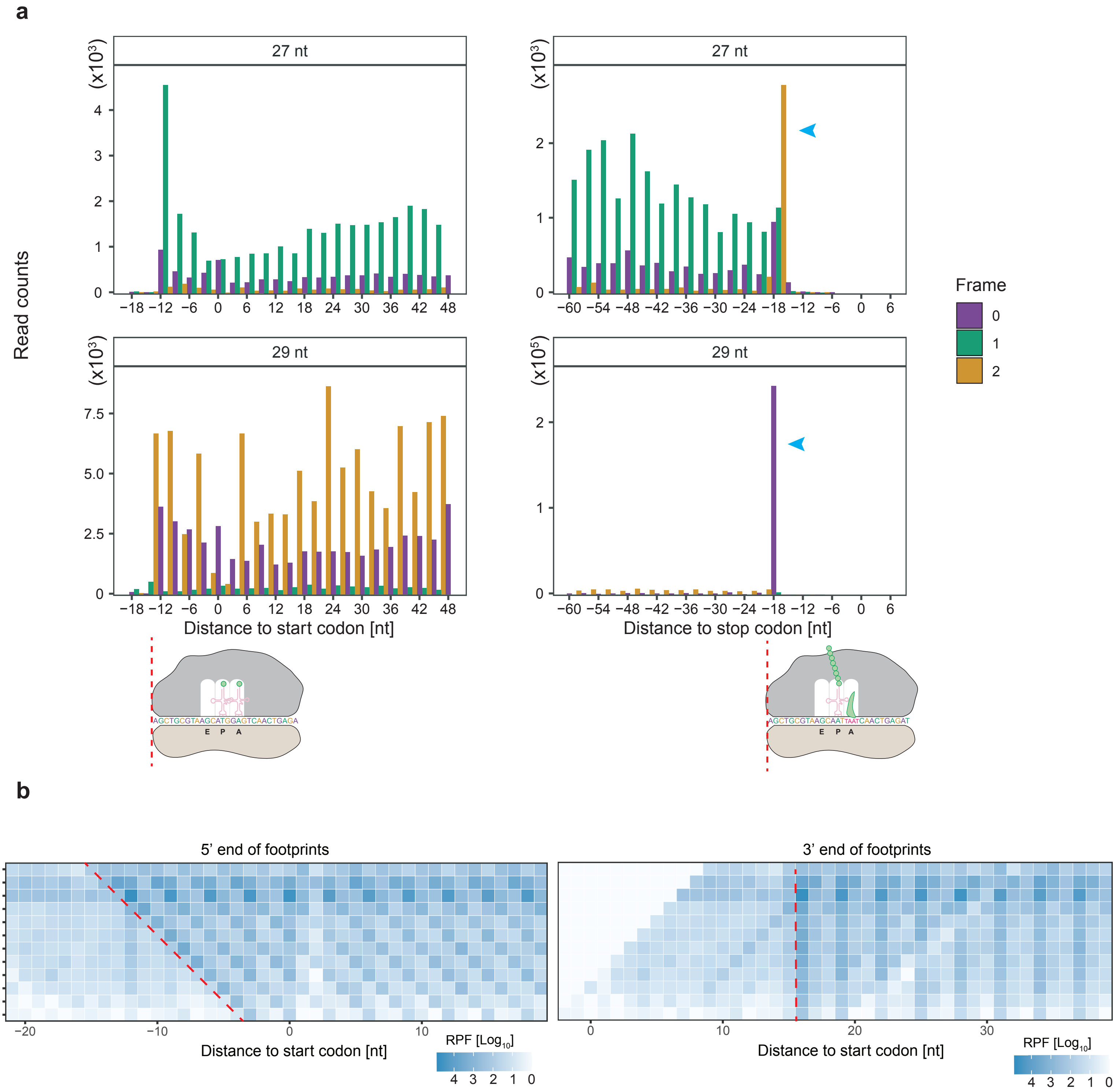


**Supplementary Figure 1**

c

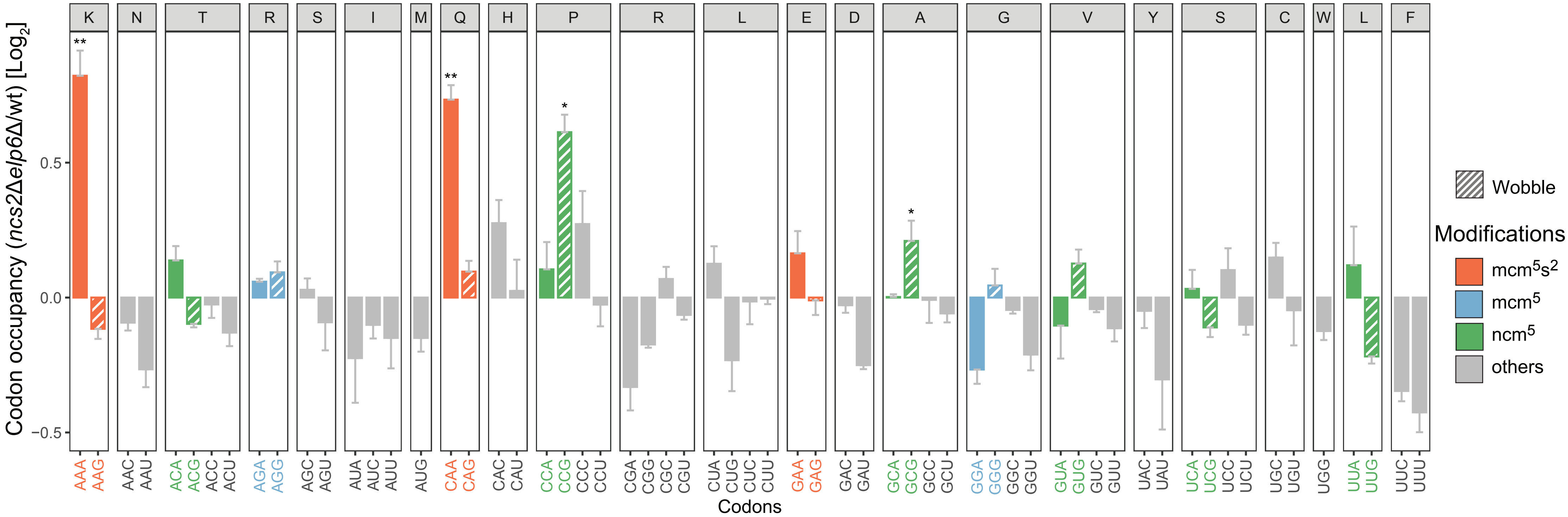


d

| Codon | Anticodon | Amino acid | Codon | Anticodon | Amino acid | Codon | Anticodon | Amino acid | Codon | Anticodon | Amino acid |
|---|---|---|---|---|---|---|---|---|---|---|---|
| UUU | — | Phe (F) | UCU | IGA (11) | Ser (S) | UAU | — | Tyr (Y) | UGU | — | Cys (R) |
| UUC | GmAA (10) | | UCC | — | | UAC | GΨA (8) | | UGC | GCA (4) | |
| UUA | ncm$^5$UmAA (7) | Leu (L) | UCA | ncm$^5$UGA (3) | | UAA | — | n.a. | UGA | — | n.a. |
| UUG | m$^5$CAA (10) | | UCG | CGA (1) | | UAG | — | | UGG | CmCA (6) | Trp (W) |
| CUU | — | Leu (L) | CCU | IGG (2) | Pro (P) | CAU | — | His (H) | CGU | ICG (6) | Arg (R) |
| CUC | GAG (1) | | CCC | — | | CAC | GUG (7) | | CGC | — | |
| CUA | UAG (3) | | CCA | ncm$^5$UGG (10) | | CAA | mcm$^5$s$^2$UUG (9) | Gln (Q) | CGA | — | |
| CUG | — | | CCG | — | | CAG | CUG (1) | | CGG | CCG (1) | |
| AUU | IAU (13) | Ile (I) | ACU | IGU (11) | Thr (T) | AAU | — | Asn (N) | AGU | — | Ser (S) |
| AUC | — | | ACC | — | | AAC | GUU (10) | | AGC | GCU (4) | |
| AUA | ΨAΨ(2) | | ACA | ncm$^5$UGU (4) | | AAA | mcm$^5$s$^2$UUU (7) | Lys (K) | AGA | mcm$^5$UCU (11) | Arg (R) |
| AUG | CAU (5/5) | Met (M) | ACG | CGU (1) | | AAG | CUU (14) | | AGG | CCU (1) | |
| GUU | IAC (14) | Val (V) | GCU | IGC (11) | Ala (A) | GAU | — | Asp (D) | GGU | — | Gly (G) |
| GUC | — | | GCC | — | | GAC | GUC (16) | | GGC | GCC (16) | |
| GUA | ncm$^5$UAC (2) | | GCA | ncm$^5$UGC (5) | | GAA | mcm$^5$s$^2$UUC (14) | Glu (E) | GGA | mcm$^5$UCC (3) | |
| GUG | CAC (2) | | GCG | — | | GAG | CUC (2) | | GGG | CCC (2) | |

**Supplementary Figure 1**

**Supplementary Fig. 1. $U_{34}$ modifications affect translation differentially.** (a) The distribution of the 5' ends of 27- (top) and 29-nt (bottom) footprints is shown at the start (left) or at the stop (right) codon; the cartoon below depicts initiating and terminating ribosomes. The terminating ribosomes accommodate four nucleotides in the ribosomal A site due to eRF1 recognition, which is visible as an apparent frameshift at the stop codon (blue arrowhead). (b) The 5' end of initiating ribosome footprints shifts one nucleotide at a time with the length of footprints, while the 3' end of initiating ribosome footprints does not change with the length footprints. (c) Alterations in codon occupancy in the absence of $U_{34}$ tRNA modification using a position-based normalization strategy[1]. Even though AGG, GGG, and GUG do not score as significantly as in the read-based method (**Figure 1D**), they are slow compared to their synonymous A-ending codons. (d) Codon table including the tRNA anticodon with modifications. The numbers in brackets are the copy numbers of the tRNA genes in yeast. Known essential (blue) and non-essential (orange) tRNAs are marked with stars. Red dots connected by lines depict the modification-mediated wobble pairing discovered in this study (Codon table was adapted from Ref. 2).

**a**

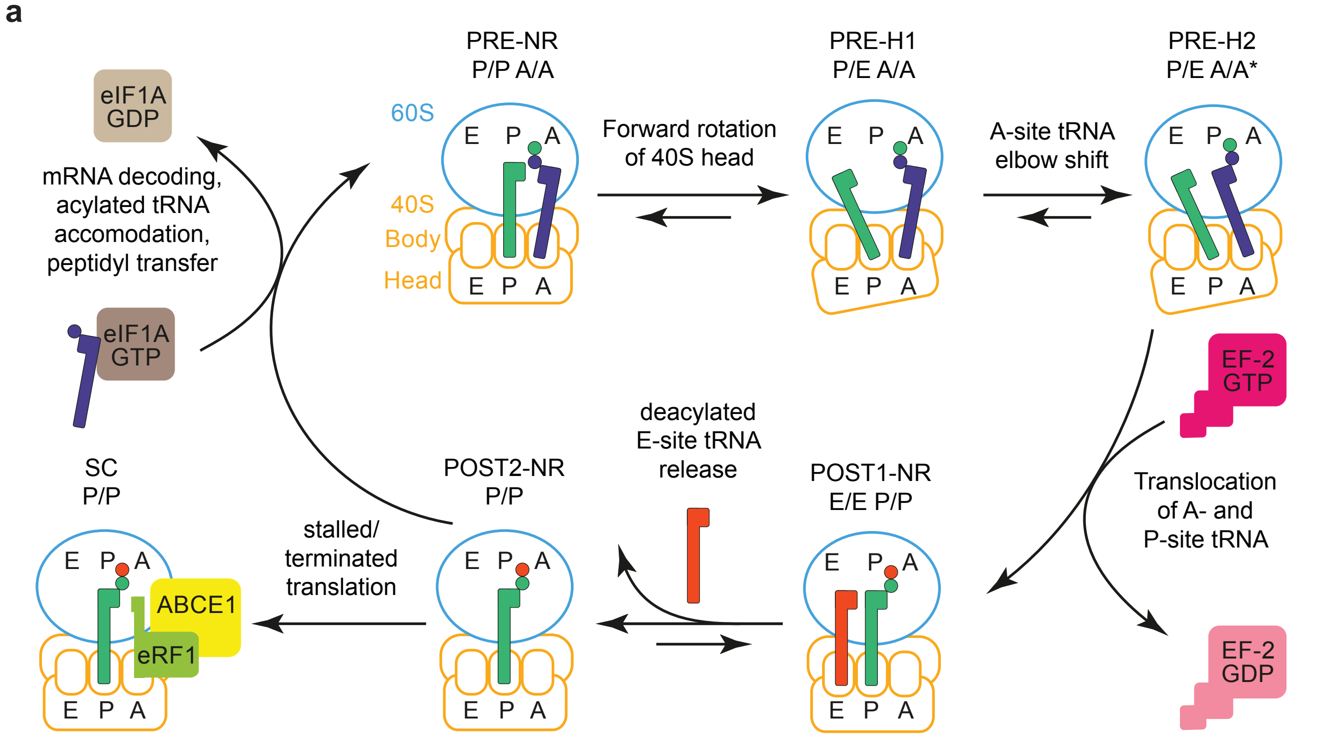


**b**

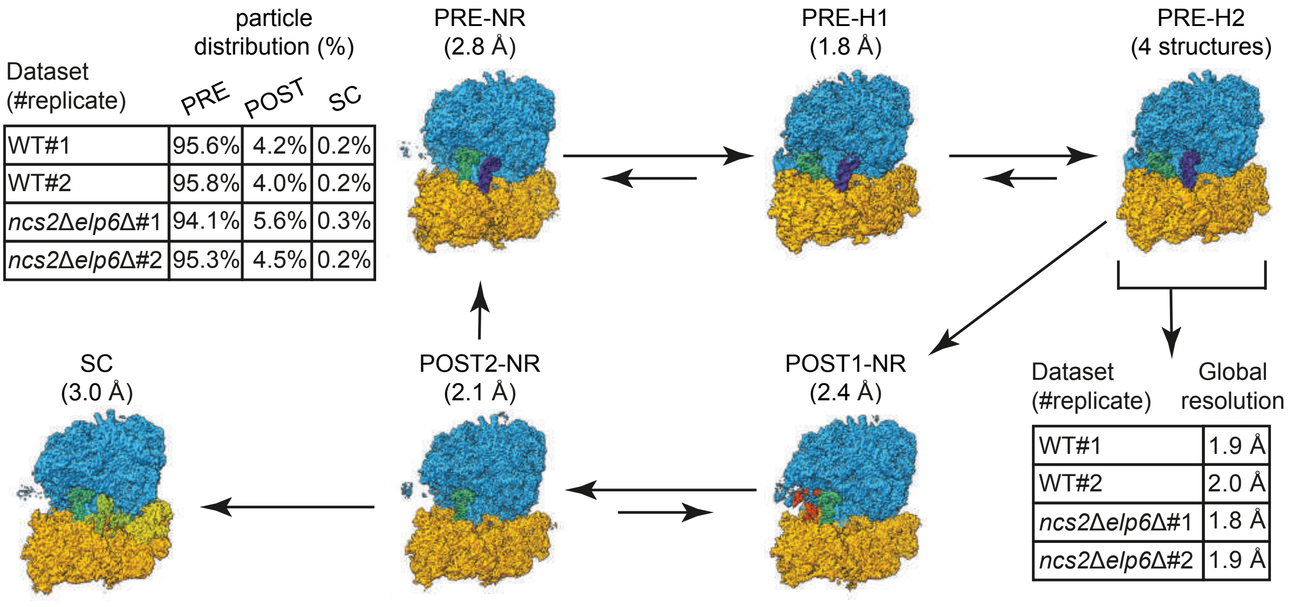


| Dataset (#replicate) | PRE | POST | SC |
|---|---|---|---|
| WT#1 | 95.6% | 4.2% | 0.2% |
| WT#2 | 95.8% | 4.0% | 0.2% |
| *ncs2Δelp6Δ*#1 | 94.1% | 5.6% | 0.3% |
| *ncs2Δelp6Δ*#2 | 95.3% | 4.5% | 0.2% |

(particle distribution (%))

| Dataset (#replicate) | Global resolution |
|---|---|
| WT#1 | 1.9 Å |
| WT#2 | 2.0 Å |
| *ncs2Δelp6Δ*#1 | 1.8 Å |
| *ncs2Δelp6Δ*#2 | 1.9 Å |

**Supplementary Figure 2**

c

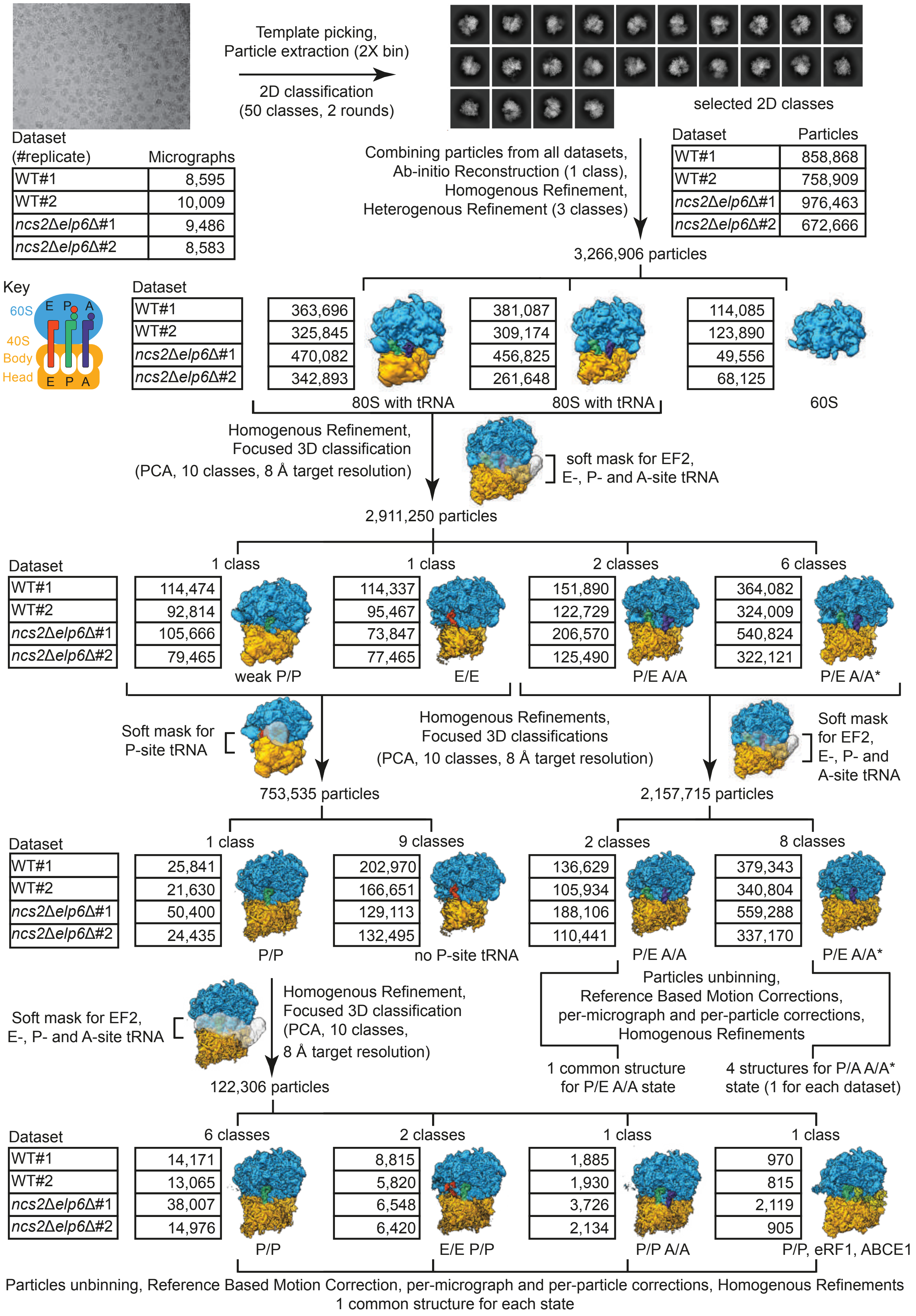


**Supplementary Figure 2**

d

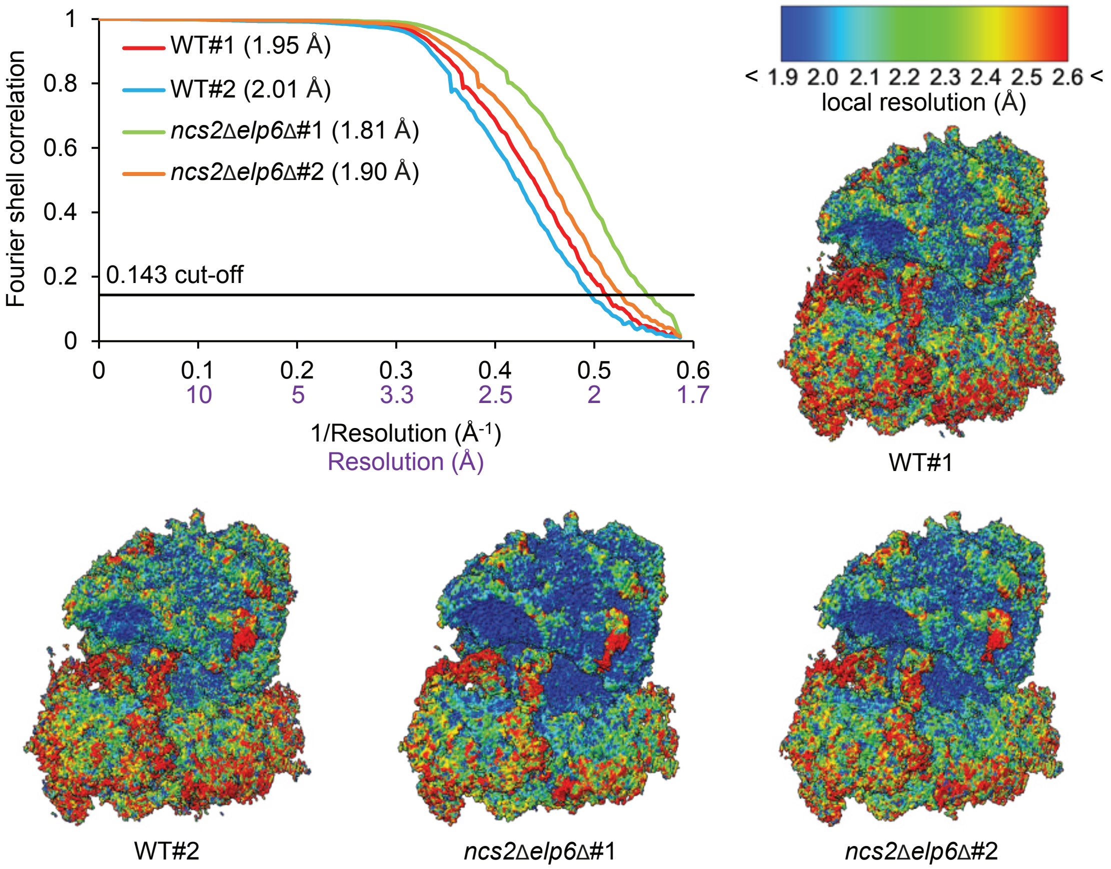


e

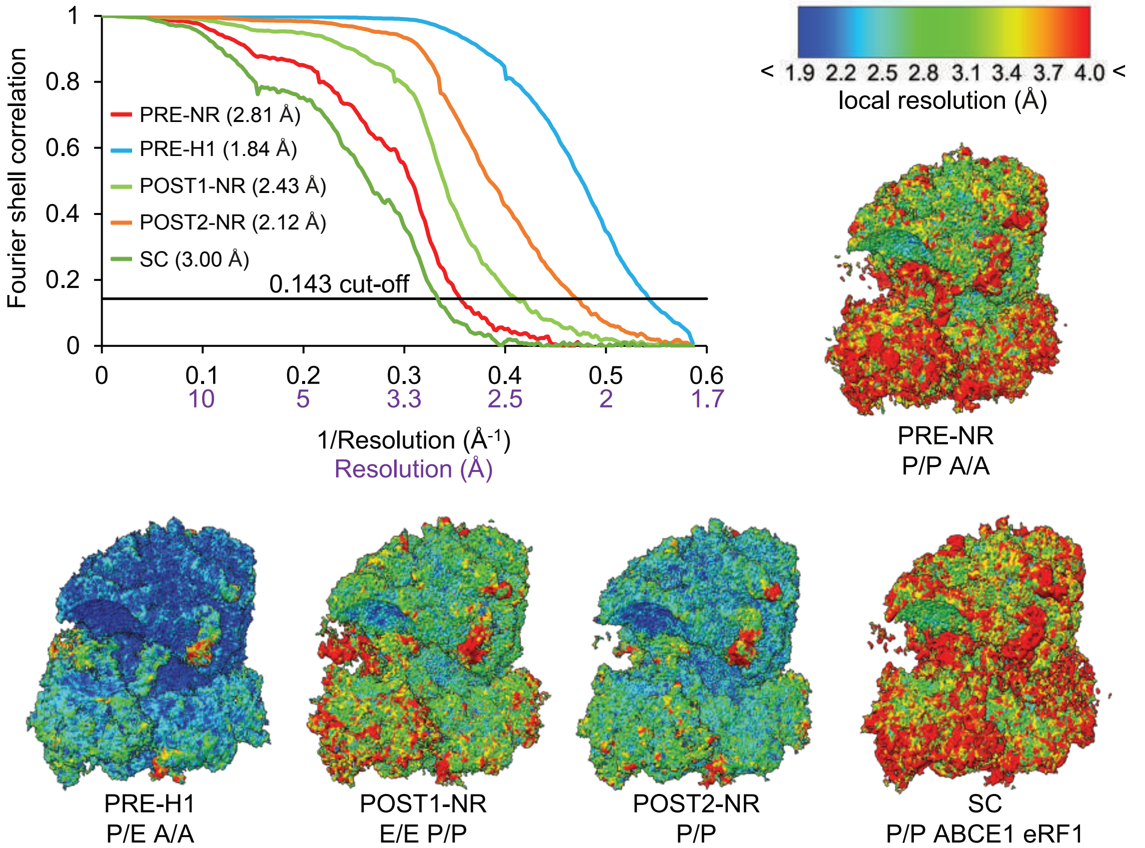


**Supplementary Figure 2**

**Supplementary Fig. 2. Cryo-EM analysis of wild-type and *ncs2*Δ*elp6*Δ ribosomes.** (a-e) Cryo-EM reconstructions of wild-type (WT) and *ncs2*Δ*elp6*Δ *S. cerevisiae* translating ribosomes extracted by MNase digestion from polysomes. (a) Schematic depiction 80S ribosomes and relative positioning of 60S (blue) and 40S subunit (orange), E-, P-, and A-site tRNA (red, dark green, and purple) during translation elongation, before (PRE) and after translocation (POST). The forward and backward arrows indicate the transition and potential reversibility of intermediates. Six 80S translation elongation states reported in the current study are as follows: PRE-translocation non-rotated P/P A/A (PRE-NR); PRE-translocation-hybrid P/E A/A (PRE-H1); PRE-translocation-hybrid P/E A/A* (PRE-H2); post-translocation non-rotated E/E P/P (POST1-NR); post-translocation non-rotated P/P (POST2-NR); and splitting complex (SC) with P/P tRNA, eRF1 (light green), and ABCE1 (yellow). (b) Relative distribution (%) of 80S particles into PRE, POST and SC states and corresponding cryo-EM reconstructions. The cryo-EM datasets were collected in 2 replicates (#1/#2) for each WT and *ncs2*Δ*elp6*Δ *S. cerevisiae* strain. Global resolutions (in Å) are reported for each cryo-EM structure (GSFSC with 0.143 cut-off). (c) Classification scheme of cryo-EM data for bulk 80S ribosomes from wild-type and *ncs2*Δ*elp6*Δ yeast. The diagram depicts the strategy applied to remove junk particles by 2D classification and to separate the ribosomal particles into different states by Heterogenous Refinement and 3D classification. The Heterogenous Refinement allowed the removal of particles corresponding to the 60S subunit. The focused 3D classifications allowed the removal of the particles corresponding to non-translating 80S ribosomes (lacking P-site tRNA). Furthermore, the approach allowed separation and reconstruction of the following translationally-active 80S ribosomes: PRE-translocation non-rotated P/P A/A (PRE-NR); PRE-translocation-hybrid P/E A/A (PRE-H1); PRE-translocation-hybrid P/E A/A* (PRE-H2); post-translocation non-rotated E/E P/P (POST1-NR); post-translocation non-rotated P/P (POST2-NR); and splitting complex with P/P tRNA, eRF1, and ABCE1 (SC). (d) Global and local resolution estimations of cryo-EM maps for 80S ribosome PRE-H2 state with A/P A/A* tRNA. Fourier shell correlation (FSC) for cryo-EM maps of ribosomes, from two independent isolations (#1/#2), from wild-type (WT) or *ncs2*Δ*elp6*Δ yeast. The global and local resolution indicated for final maps was estimated using a 0.143 FSC cut-off. (e) Global and local resolution estimations of cryo-EM maps for 80S ribosomes with P-site tRNA. Fourier shell correlation (FSC) for cryo-EM maps of ribosomes in the following states: PRE-translocation non-rotated P/P A/A (PRE-NR); PRE-translocation-hybrid P/E A/A (PRE-H1); post-translocation non-rotated E/E P/P (POST1-NR); post-translocation non-rotated P/P (POST2-NR); and splitting complex (SC) with P/P tRNA, eRF1, and ABCE1. The global and local resolutions indicated for final maps was estimated using a 0.143 FSC cut-off.

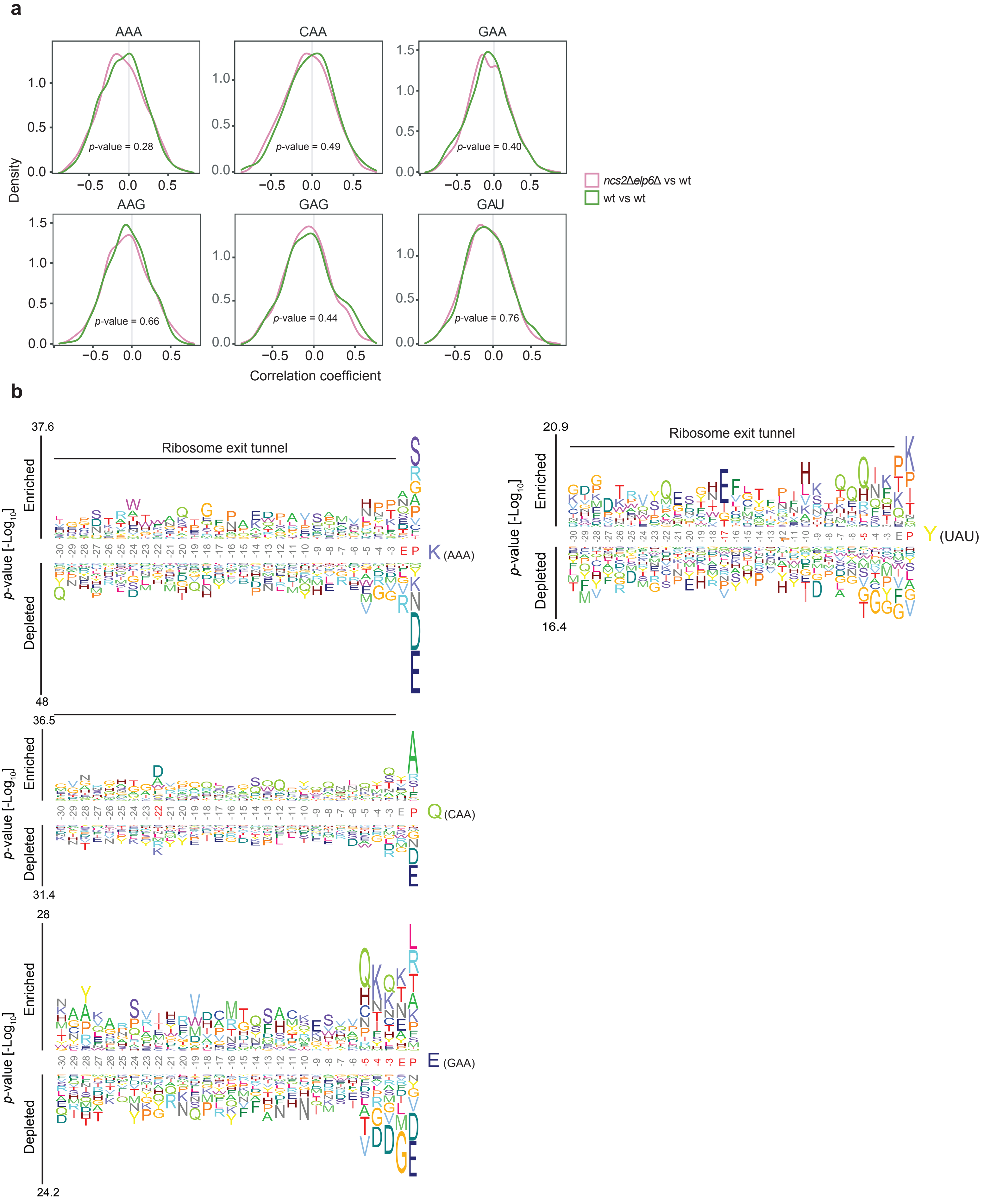


Supplementary Figure 3

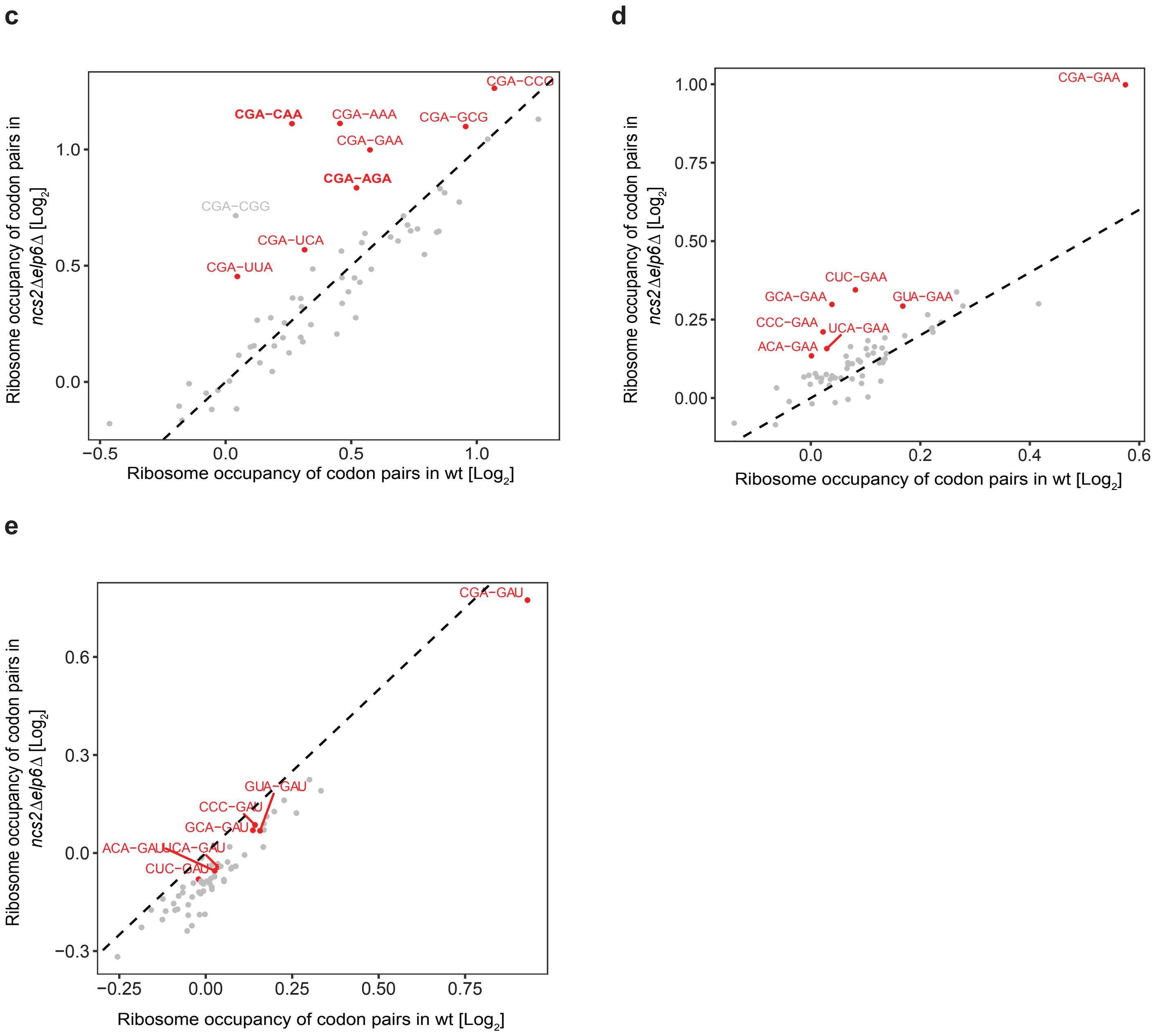


**Supplementary Figure 3**

**Supplementary Fig. 3. Strong pausing occurs at tRNA modification-dependent codon pairs.** (a) Correlation analyses between *ncs2Δelp6Δ* and wild type was performed using vulnerability scores against relative positions of each codon in each ORF. For comparison, the same analysis was performed using two wild-type replicates (two-sided Kolmogorov-Smirnov test. *p*-values are indicated). (b) Amino acid motifs upstream of AAA (top left), CAA (middle left), GAA (bottom left) and UAU (right) codons (~30 codons in the ribosome exit tunnel) with high vulnerability scores (top 1000 codons are selected). Red letters or numbers indicate positions in the motif that are statistically significant (Bonferroni-corrected *p*-value < 0.01). (c-e) Relative ribosome occupancy of specific codon pairs between *ncs2Δelp6Δ* and wild-type yeast. CGA-NNN (c) and NNN-GAA (d) codon pairs, and NNN-GAU (e) codon pairs that are not affected by $U_{34}$ tRNA modifications are plotted. $U_{34}$ modification-dependent codon pairs are highlighted in red in (c) and (d) but they are not slow in (e).

**a**

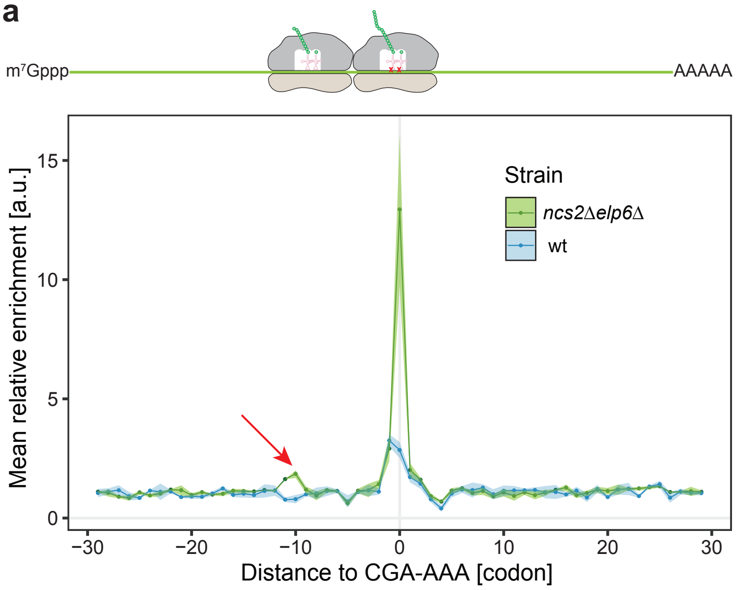


**b**

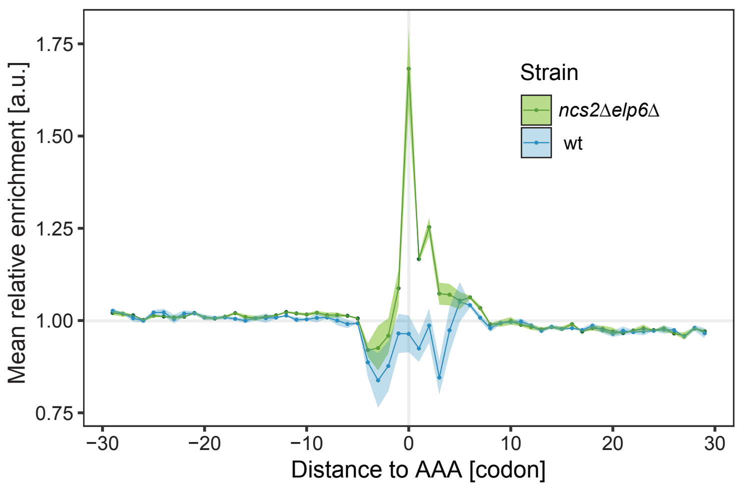


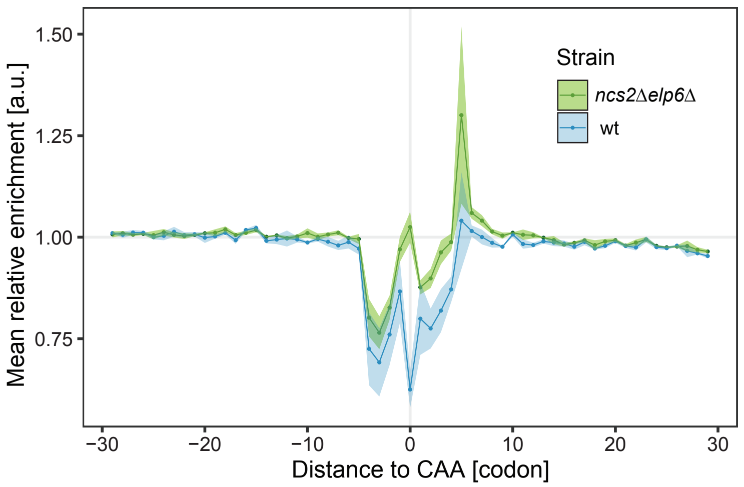


**Supplementary Figure 4**

**Supplementary Fig. 4. Ribosomes stall on slow codon pairs.** (a) and (b) The distribution of ribosome occupancy (monosomes) around CGA-AAA (a), AAA (b, left), and CAA (b, right). Colliding ribosomes ~10 codons upstream of the codon pairs in the *ncs2*Δ*elp6*Δ mutant are highlighted by a red arrow. The shaded area indicates the degree of experimental variation within three replicates. The cartoon above (a) depicts collided ribosomes relative to the plot. Note that no queueing ribosomes are observed upstream of individual AAA and CAA codons.

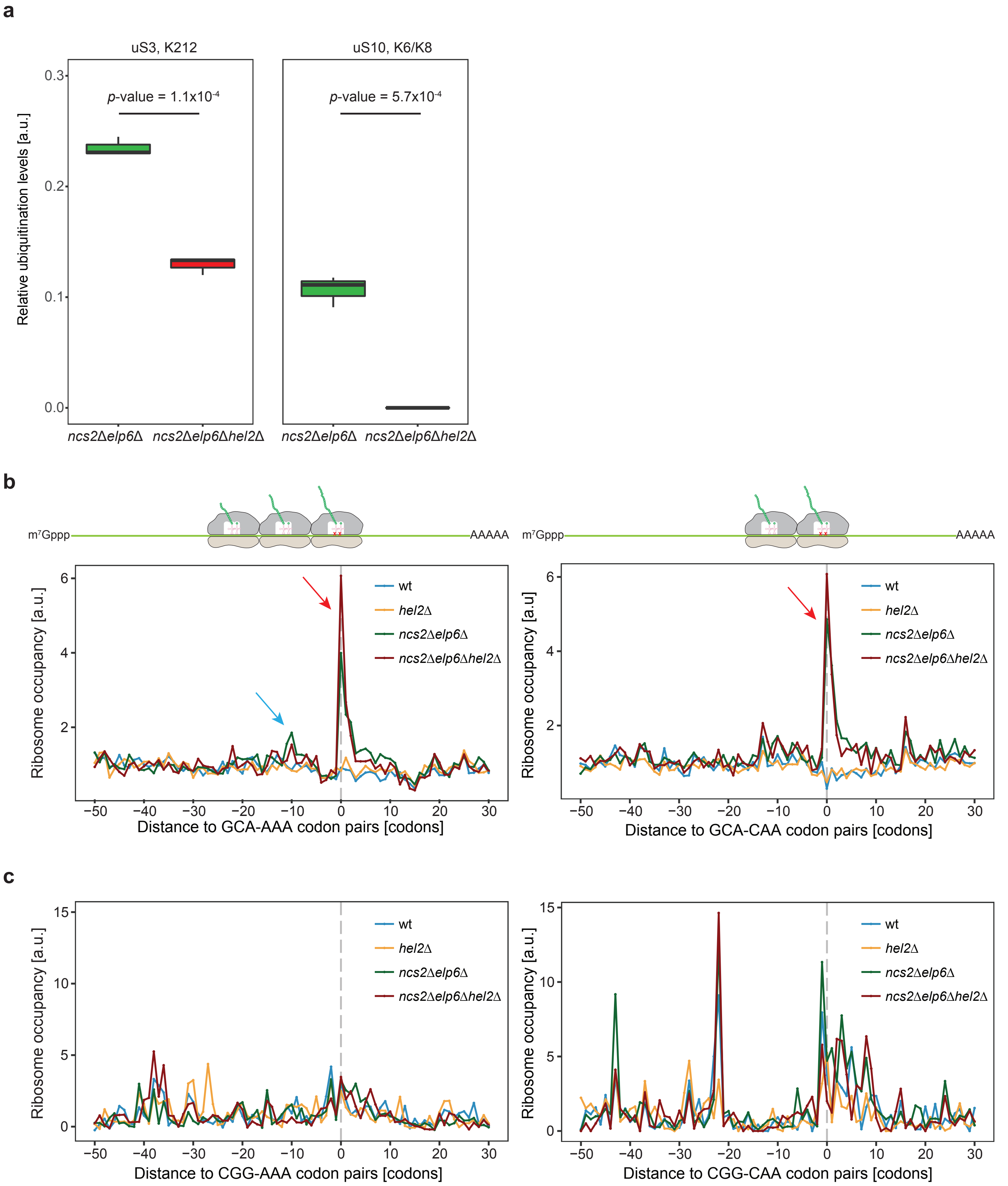


**Supplementary Figure 5**

**d**

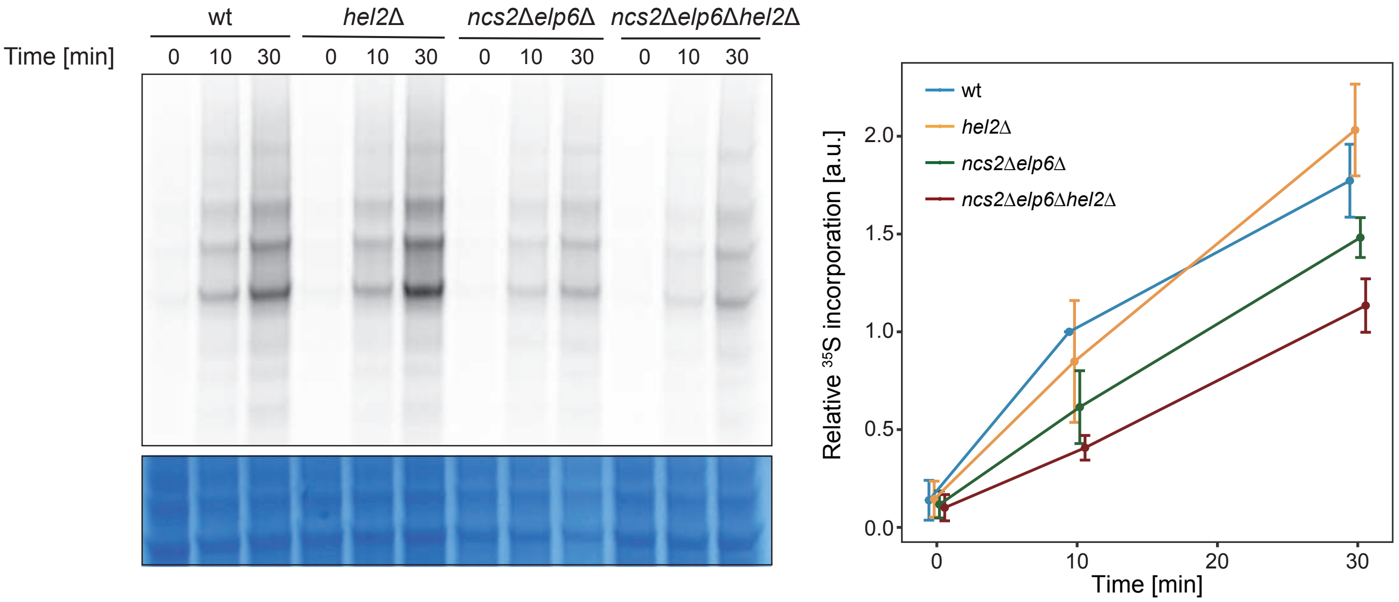


**e**

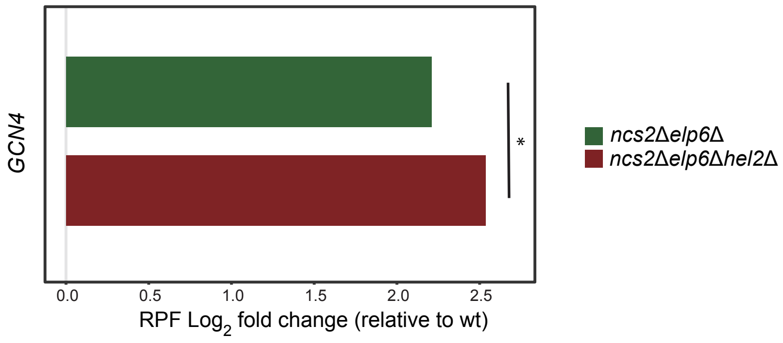


**f**

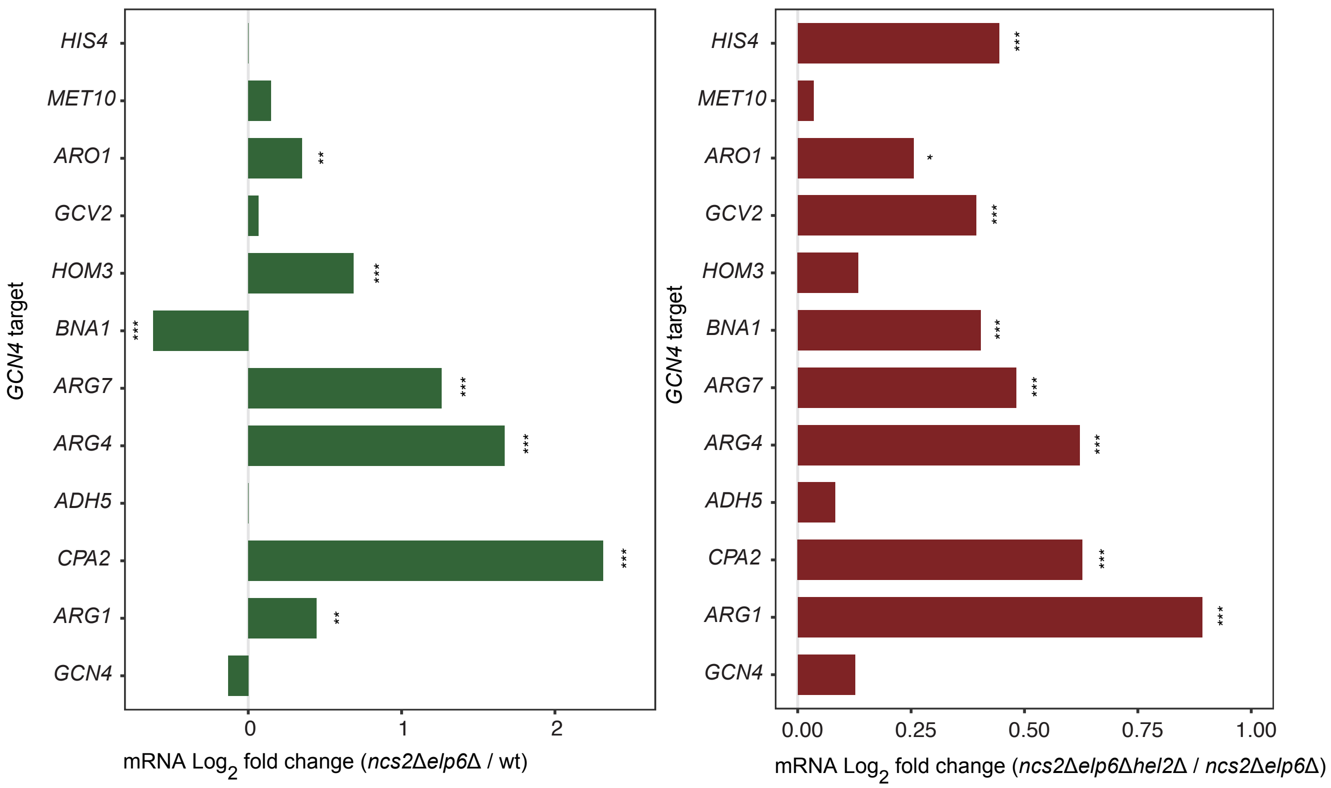


**Supplementary Figure 5**

**g**

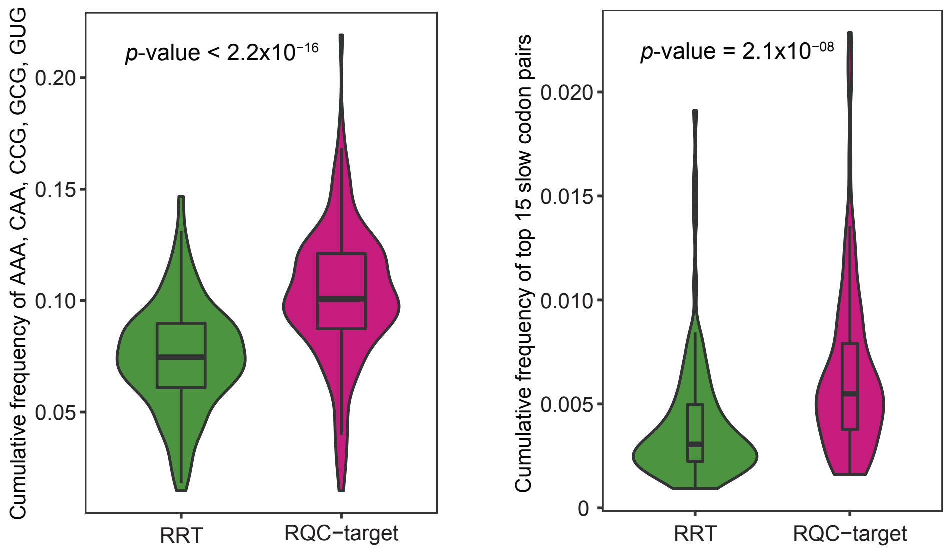


**h**

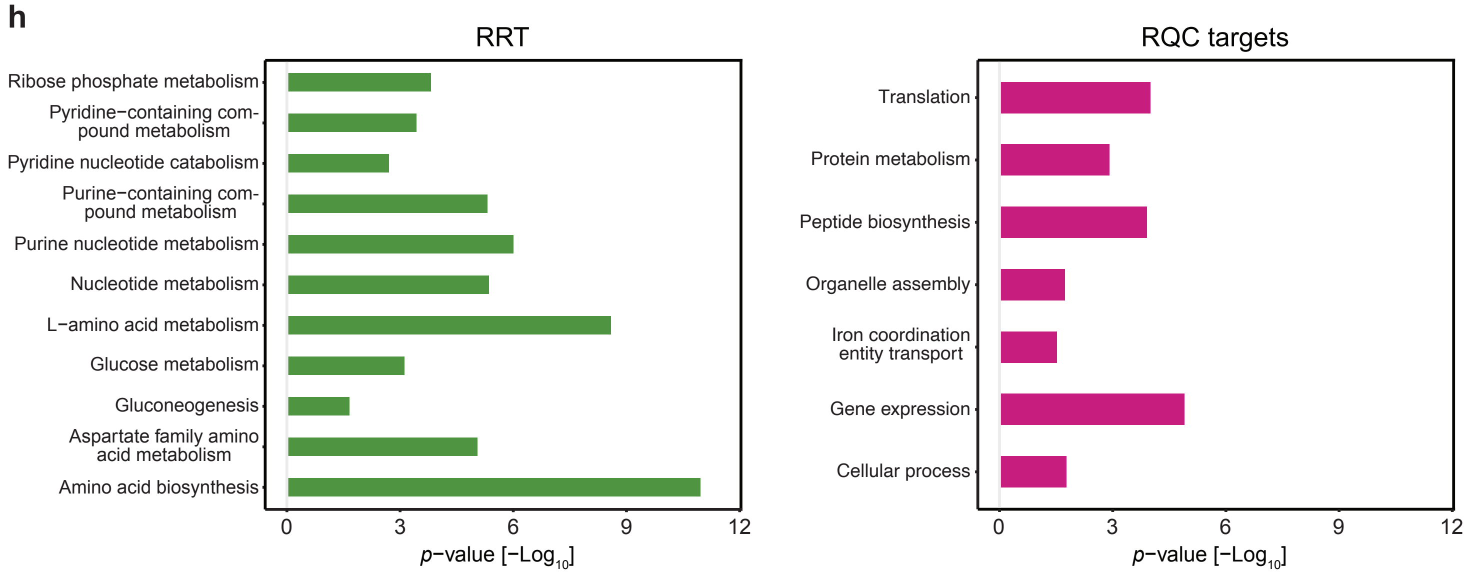


**i**

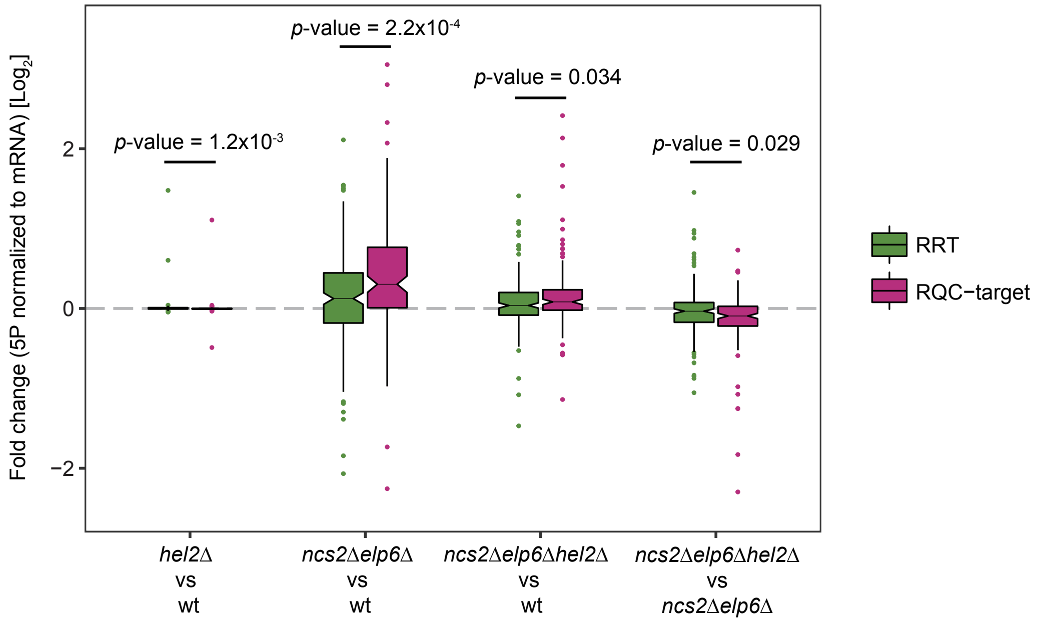


**Supplementary Figure 5**

**Supplementary Fig. 5. Ribosomes collide and are targeted by ribosome-associated quality control (RQC).** (a) Quantification of ubiquitination levels of uS3 and uS10 by mass spectrometry in the *ncs2*Δ*elp6*Δ and *ncs2*Δ*elp6*Δ*hel2*Δ backgrounds (two-sided student's t-test). (b) Disome occupancy around GCA-AAA (left) and GCA-CAA (right) codon pairs in wild-type, *hel2*Δ, *ncs2*Δ*elp6*Δ and *ncs2*Δ*elp6*Δ*hel2*Δ yeast. The average disome occupancy from two replicates was plotted. Peaks at position 0 represent the A site of the first stalling ribosome (red arrow). One queuing disome (blue arrow) was observed upstream of GCA-AAA, indicating a total of three ribosomes at the stalling site. The cartoon above depicts collided ribosomes relative to the plot. (c) Disome occupancy around the control codon pairs CGG-AAA and CGG-CAA in the same yeast strains. (d) Metabolic labeling of newly synthesized proteins after 0, 10 or 30 min incubation with $^{35}$S L-methionine in wild-type, *hel2*Δ, *ncs2*Δ*elp6*Δ and *ncs2*Δ*elp6*Δ*hel2*Δ cells. Translation is strikingly impaired in *ncs2*Δ*elp6*Δ*hel2*Δ compared to *ncs2*Δ*elp6*Δ. A representative gel is shown on the left and quantification of three replicates is shown on the right. (e) Expression levels (RPF) of *GCN4* in *ncs2*Δ*elp6*Δ and *ncs2*Δ*elp6*Δ*hel2*Δ compared to wild type. (f) The transcription of Gcn4-targets was mostly upregulated in *ncs2*Δ*elp6*Δ yeast compared to wild type (left), and the upregulation is further increased in *ncs2*Δ*elp6*Δ*hel2*Δ (right). However, the transcription of *GCN4* itself does not change, indicating that the integrated stress response (ISR) is activated in the absence of ribosome-associated quality control (RQC) (two-sided student's t-test; *p-value < 0.05, **p-value < 0.01; ***p-value < 0.001). (g) The frequency of slow codons (left) or slow codon pairs (right) was compared between RQC-targets and RQC-refractory transcripts (RRTs) (one-sided Mann-Whitney U-test). (h) Gene ontology analysis of RRT (left) and RQC-targets (right). Significant terms in biological process are shown (Fisher's exact test). (i) Differential mRNA degradation analysis by 5PSeq in wild-type, *hel2*Δ, *ncs2*Δ*elp6*Δ and *ncs2*Δ*elp6*Δ*hel2*Δ cells. The analysis was performed by normalizing to mRNA level to exclude transcriptional effects. The degradation of RQC-target mRNAs was rescued in *ncs2*Δ*elp6*Δ*hel2*Δ compared to *ncs2*Δ*elp6*Δ yeast (two-sided Mann-Whitney U-test).

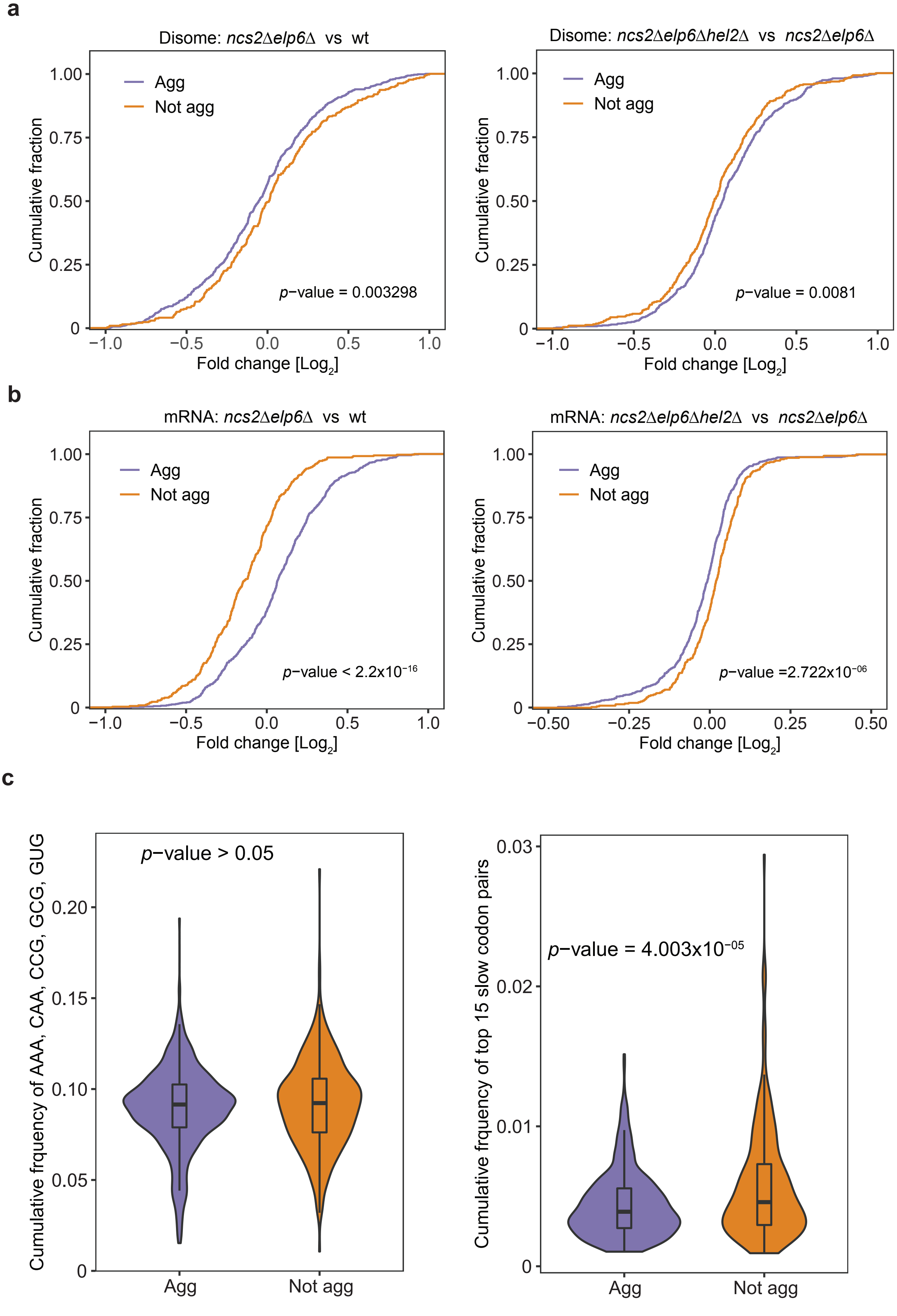


**Supplementary Figure 6**

**d**

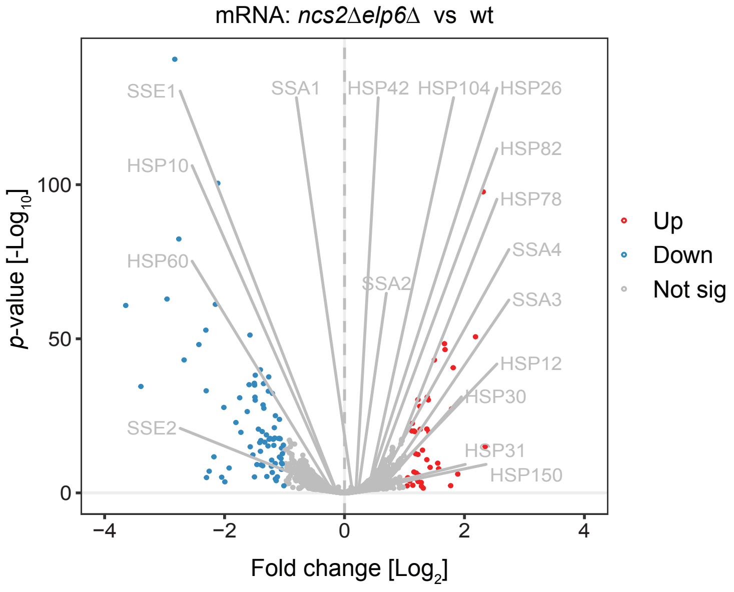


**e**

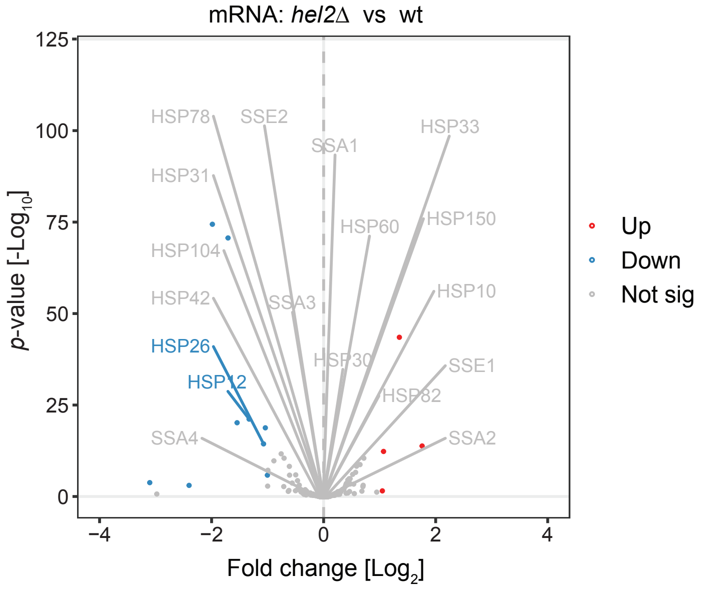


**f**

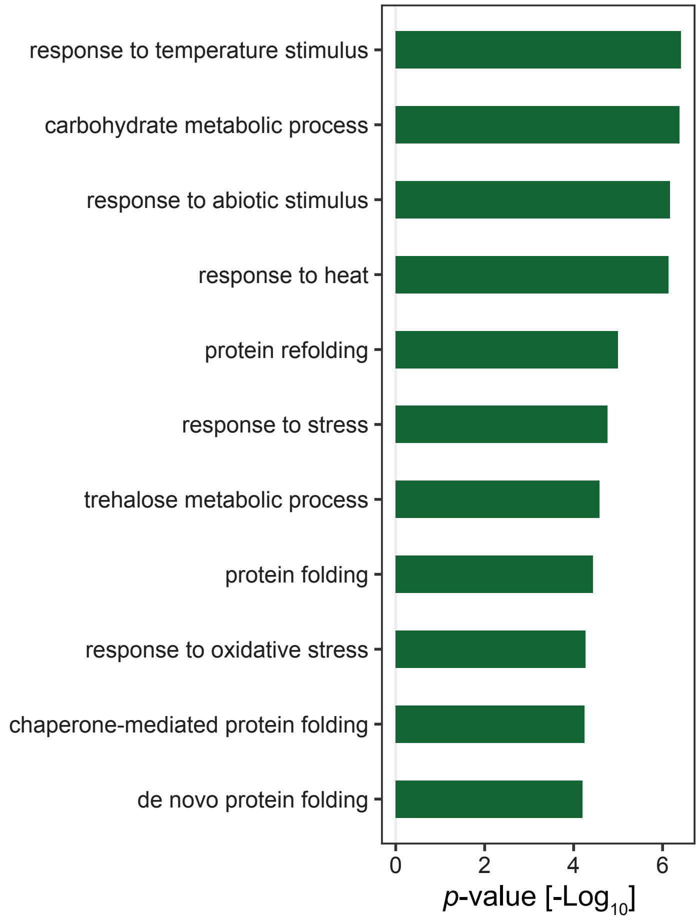


**Supplementary Figure 6**

**Supplementary Fig. 6. The RQC pathway alleviates protein homeostasis defects in coordination with chaperones.** (a) The disome occupancy was compared between *ncs2*Δ*elp6*Δ and wild type (left) or between *ncs2*Δ*elp6*Δ*hel2*Δ and *ncs2*Δ*elp6*Δ (right) for aggregated (n=610) and non-aggregating (n=407) proteins using DESeq2[3]. Compared to aggregated proteins, disomes are more abundant on transcripts that encode non-aggregating proteins in the absence of $U_{34}$ tRNA modifications (left) but are less abundant on these transcripts when *HEL2* is additionally deleted (right; one-sided Kolmogorov-Smirnov test). Monosome levels were used for normalization like for Fig. 4b. (b) mRNA levels of non-aggregating proteins are lower in *ncs2*Δ*elp6*Δ yeast (left) and this effect is rescued in *ncs2*Δ*elp6*Δ*hel2*Δ cells (right) (one-sided Kolmogorov-Smirnov test). (c) The frequency of $U_{34}$ modification-related slow codons (left) or codon pairs (right) were compared between aggregated proteins and non-aggregating proteins (one-sided Mann-Whitney U-test). (d) and (e) DESeq2 differential expression analysis of mRNA levels comparing *ncs2*Δ*elp6*Δ (d) and *hel2*Δ (e) to wild type, respectively. Chaperones are generally not altered in these two mutant strains. Up- or downregulated genes with $p$-adjusted value < 0.05 and $Log_2$ fold change > 1 are highlighted in red or blue. (f) Gene ontology analysis of genes that are upregulated in *ncs2*Δ*elp6*Δ*hel2*Δ compared to *ncs2*Δ*elp6*Δ. Significant terms in biological process are shown (Fisher's exact test).
